# Comprehensive epigenome characterization reveals diverse transcriptional regulation across human vascular endothelial cells

**DOI:** 10.1101/756056

**Authors:** Ryuichiro Nakato, Youichiro Wada, Ryo Nakaki, Genta Nagae, Yuki Katou, Shuichi Tsutsumi, Natsu Nakajima, Hiroshi Fukuhara, Atsushi Iguchi, Takahide Kohro, Yasuharu Kanki, Yutaka Saito, Mika Kobayashi, Akashi Izumi-Taguchi, Naoki Osato, Kenji Tatsuno, Asuka Kamio, Yoko Hayashi-Takanaka, Hiromi Wada, Shinzo Ohta, Masanori Aikawa, Hiroyuki Nakajima, Masaki Nakamura, Rebecca C. McGee, Kyle W. Heppner, Tatsuo Kawakatsu, Michiru Genno, Hiroshi Yanase, Haruki Kume, Takaaki Senbonmatsu, Yukio Homma, Shigeyuki Nishimura, Toutai Mitsuyama, Hiroyuki Aburatani, Hiroshi Kimura, Katsuhiko Shirahige

## Abstract

**Background:** Endothelial cells (ECs) make up the innermost layer throughout the entire vasculature. Their phenotypes and physiological functions are initially regulated by developmental signals and extracellular stimuli. The underlying molecular mechanisms responsible for the diverse phenotypes of ECs from different organs are not well understood.

**Results:** To characterize the transcriptomic and epigenomic landscape in the vascular system, we cataloged gene expression and active histone marks in nine types of human ECs (generating 148 genome-wide datasets) and carried out a comprehensive analysis with chromatin interaction data. We identified 3,765 EC-specific enhancers, some of which were associated with disease-associated genetic variations. We also identified various candidate marker genes for each EC type. Notably, reflecting the developmental origins of ECs and their roles in angiogenesis, vasculogenesis and wound healing.

**Conclusions:** While the importance of several HOX genes for early vascular development and adult angiogenesis in pathological conditions has been reported, a systematic analysis of the regulation and roles of HOX genes in mature tissue cells has been lacking. These datasets provide a valuable resource for understanding the vascular system and associated diseases.

## INTRODUCTION

Endothelial cells (ECs), which make up the innermost blood vessel lining of the body, express diverse phenotypes that affect their morphology, physiological function and gene expression patterns in response to the extracellular environment, including the oxygen concentration, blood pressure and physiological stress. In the kidney, for example, the vascular bed plays a role in the filtration of blood; in the brain, however, the vascular architecture protects the central nervous system from toxins and other components of the blood (1). Endothelial heterogeneity is mainly dependent on both the function of each organ and the developmental lineage of different EC populations, which result in adaptation to the vascular microenvironment. It is widely recognized that certain specific vessels are susceptible to pathological changes, which include those related to atherosclerosis and inflammation (2). Atherosclerosis, which occurs in the muscular and elastic arteries, is a progressive disease characterized by the accumulation of macrophages, and this process is initiated by the expression of cell adhesion molecules, such as P-selectin. In mice, the expression level of P-selectin is higher in the lung and mesentery vesicles compared with the heart, brain, stomach and muscle (3).

In clinical practice, the thoracic, radial and gastroepiploic arteries are used for coronary bypass grafts because these arteries have no tendency toward atherosclerosis and hence are therapeutically advantageous in patients with coronary artery plaques (4). In addition, the long-lasting results from coronary bypass graft surgery indicate that vessels transplanted to a new environment differ in their outcome based on their origin as an artery or vein (5, 6). Although the elucidation of the molecular mechanisms underlying EC heterogeneity is critically important for the development of vascular bed–specific remedies, these mechanisms have remained largely unknown because ECs do not display such heterogeneity when cultured *in vitro*.

Epigenetic variation is a prime candidate for controlling the heterogeneity among various ECs. Increasing evidence supports the idea that certain site-specific characteristics are epigenetically regulated and easily altered by changes in the human extracellular microenvironment. Previous gene expression studies of many types of human ECs in culture demonstrated that site-specific epigenetic modifications play an important role in differential gene expression (7). Moreover, our recent reports elucidated that there are different histone modifications present in the same genomic loci, such as *GATA6*, in human umbilical vein endothelial cells (HUVECs) and human dermal microvascular endothelial cells (HMVECs) (8, 9). Despite the discovery of these important insights, we still lack a systematic understanding of how the epigenomic landscape contributes to EC phenotype and heterogeneity. Therefore, there is a great demand for a comprehensive epigenomic catalog of the various EC types.

As a part of the International Human Epigenome Consortium (IHEC) project (10), we collected Chromatin immunoprecipitation followed by sequencing (ChIP-seq) data for the active histone modifications trimethylated H3 at Lys4 (H3K4me3) and acetylated H3 at Lys27 (H3K27ac) in EC DNA from nine different vascular cell types from multiple donors. We implemented large-scale comparative ChIP-seq analysis of these datasets to understand how the diverse phenotypes of ECs are regulated by key genes. All datasets used in this study are publicly available and are summarized on our website (https://rnakato.github.io/HumanEndothelialEpigenome/).

## RESULTS

### Reference epigenome generation across EC types

To establish an epigenetic catalog for different EC types, we generated a total of 491 genome-wide datasets, consisting of 424 histone modification ChIP-seq and 67 paired-end RNA sequencing (RNA-seq) datasets, encompassing a total of 22.3 billion sequenced reads. ECs were maintained as primary cultures with a physiological concentration of vascular endothelial growth factor (VEGF) and a minimal number of passages (fewer than six). We generated genome-wide normalized coverage tracks and peaks for ChIP-seq data and estimated normalized gene expression values for RNA-seq data.

In this study, we selected a subset of 33 EC samples (131 datasets) as a representative set comprising nine types of vessels from the human body (Figure 1A):

- Human aortic endothelial cells (HAoECs),
- Human coronary artery endothelial cells (HCoAECs),
- Human endocardial cells (HENDCs),
- Human pulmonary artery endothelial cells (HPAECs),
- Human umbilical vein endothelial cells (HUVECs),
- Human umbilical artery endothelial cells (HUAECs),
- Human common carotid artery endothelial cells (HCCaECs),
- Human renal artery endothelial cells (HRAECs),
- Human great saphenous vein endothelial cells (HGSVECs).

**Figure 1.**
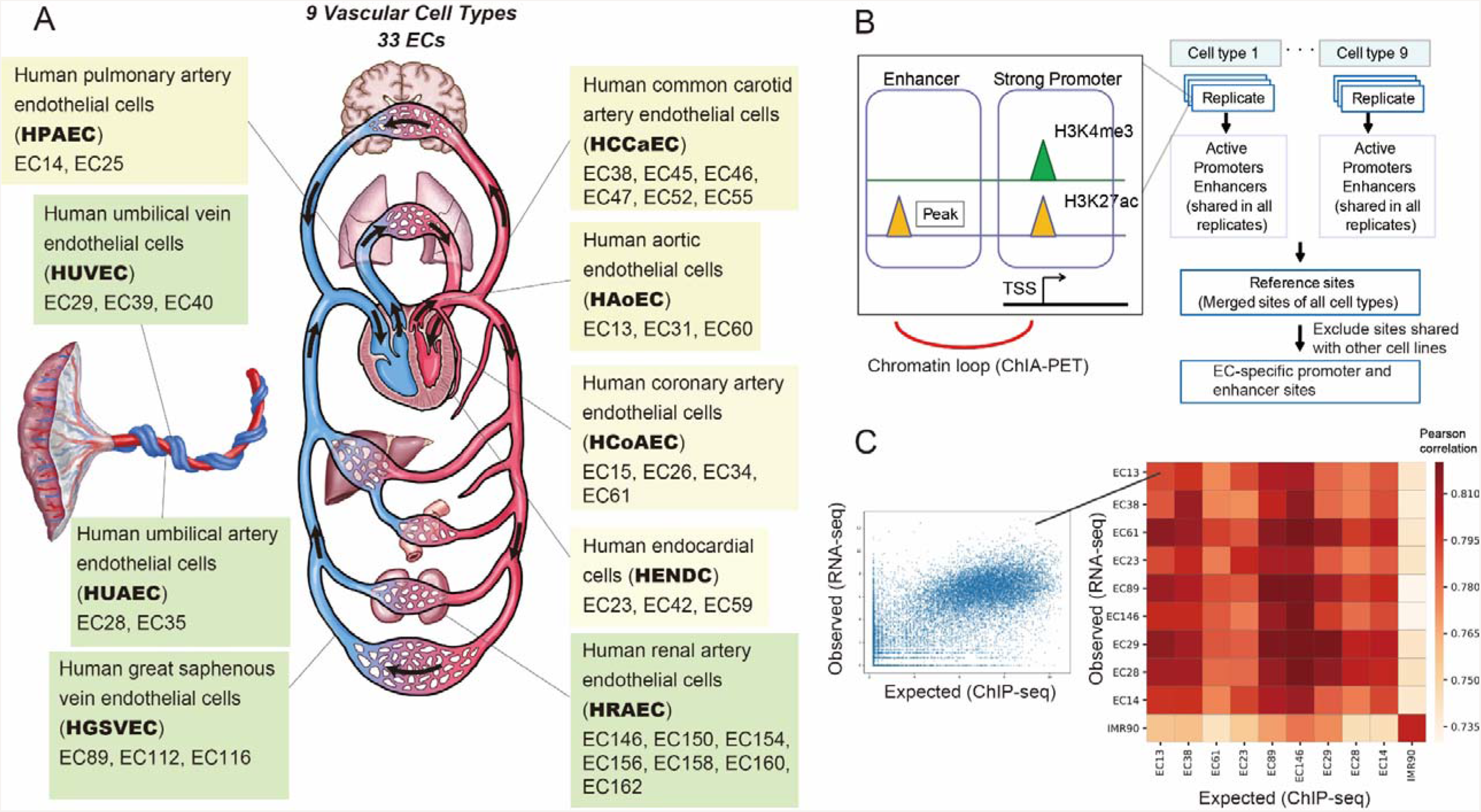
Summary of the cell types and histone modifications analyzed in this project. (A) Schematic illustration of the cardiovascular system, nine EC types and 33 individual samples (prefixed EC*) used in this paper. The yellow and green boxes indicate EC types from upper body and lower body, respectively. (B) Workflow to identify the reference sites for ECs. The active promoter and enhancer sites of each sample were identified. For each cell type, the shared sites across all samples were extracted as the reference sites. These were integrated into a single set of reference sites for ECs, which was used for the downstream analyses. ChIA-PET data were utilized to identify the corresponding gene for the reference enhancer sites. (C) Correlation between observed and expected (from ChIP-seq analysis using linear regression model) gene expression data. Left: example scatterplot of observed and expected gene expression level for genes (data from EC13). Right: Pearson correlation heatmap for representative samples of nine cell types and IMR90 cells (as a negative control).

The detail of 33 EC samples is summarized in Supplementary Table S1.

Among the structures lined by these nine EC types, a group of two aortic, six common carotid and three coronary arteries is known as the “systemic arteries” and harbors arterial blood with 100 mmHg of oxygen tension and blood pressure in a range from 140 mmHg to 60 mmHg. Data sets for each cell type comprise samples from multiple donors, all of which achieved high-quality values as evaluated below. Here we focused on two histone modifications, H3K4me3 and H3K27ac (Figure 1B), which are the key markers of active promoters and enhancers (11). Because both H3K4me3 and H3K27ac exhibit strong, sharp peaks with ChIP-seq analysis, they are more suitable for identifying shared and/or unique features across EC cell types as compared with other histone modifications that show broad peaks, such as H3K9me3.

### Quality validation

To evaluate the quality of obtained ChIP-seq data, we computed a variety of quality control measures (Supplementary Table S2), including the number of uniquely mapped reads, library complexity (the fraction of nonredundant reads), GC-content of mapped reads, genome coverage (the fraction of overlapped genomic areas with at least one mapped read), the number of peaks, signal-to-noise ratio (S/N) by the normalized strand coefficient (12), read-distribution bias measured by background uniformity (12), inter-sample correlation for each EC type and genome-wide correlation of read density across all-by-all pairs (Supplementary Figure S1). In addition, the peak distribution around several known positive/negative marker genes was visually inspected. Low-quality datasets were not used for further analyses.

To further validate the reliability of our data, we evaluated the consistency between the obtained peaks from ChIP-seq and the gene expression values from corresponding RNA-seq data. We applied a bivariate regression model (13) to estimate the expression level of all genes based on H3K4me3 and H3K27ac peaks and then calculated the Pearson correlation between the estimated and the observed expression levels from ChIP-seq and RNA-seq, respectively. We used data derived from IMR90 fibroblasts analyzed with the same antibodies as a negative control, and we confirmed that peak distribution of the ChIP-seq data was highly correlated with corresponding RNA-seq data for ECs, but not with IMR90 data (Figure 1C, Supplementary Figure S2 for the full matrix). Therefore, our ChIP-seq data are likely to represent the histone modification states of ECs for annotation.

### Identification of active promoter and enhancer sites

We used H3K4me3 and H3K27ac ChIP-seq peaks to define “active promoter (H3K4me3 and H3K27ac)” and “enhancer (H3K27ac only)” sites for each sample (Figure 1B, left). Then we assembled them and defined the common sites among all samples of a given EC type as the reference sites, to avoid differences among individuals. Finally, the reference sites of all nine EC types were merged into a single reference set for ECs (Figure 1B, right). We identified 9,121 active promoter sites (peak width, 2840.8 bp on average) and 23,202 enhancer sites (peak width, 1799.4 bp on average). The averaged peak width became relatively wide due to the merging of multiple contiguous sites.

We compared the distribution of the reference sites with gene annotation information. As expected, active promoter sites were enriched in the transcription start sites (TSSs) of genes, whereas enhancer sites were more frequently dispersed in introns and intergenic regions (Supplementary Figure S3). Among the enhancers, 15,625 (67.3%) were distally located (more than 10 kbp away from the nearest TSSs). The number of enhancer sites was more varied among the nine tissue types, whereas the number of active promoter sites was comparable across the EC types (Figure 2A, upper panel). The large number of HUAEC enhancer sites is possibly due to the small number of samples (two) and a relatively small individual difference (both samples were from newborns). We also evaluated the shared ratio of promoter and enhancer sites across all EC types (Figure 2A, lower panel). We found that nearly 80% of the active promoter sites were shared among multiple EC types. In contrast, 57.7% of the enhancers were specific to up to two EC types, suggesting that their more diverse distribution across EC types relative to active promoter sites contributes to the EC type–specific regulatory activity. These observations are consistent with previous studies for other cell lines (11, 14).

**Figure 2.**
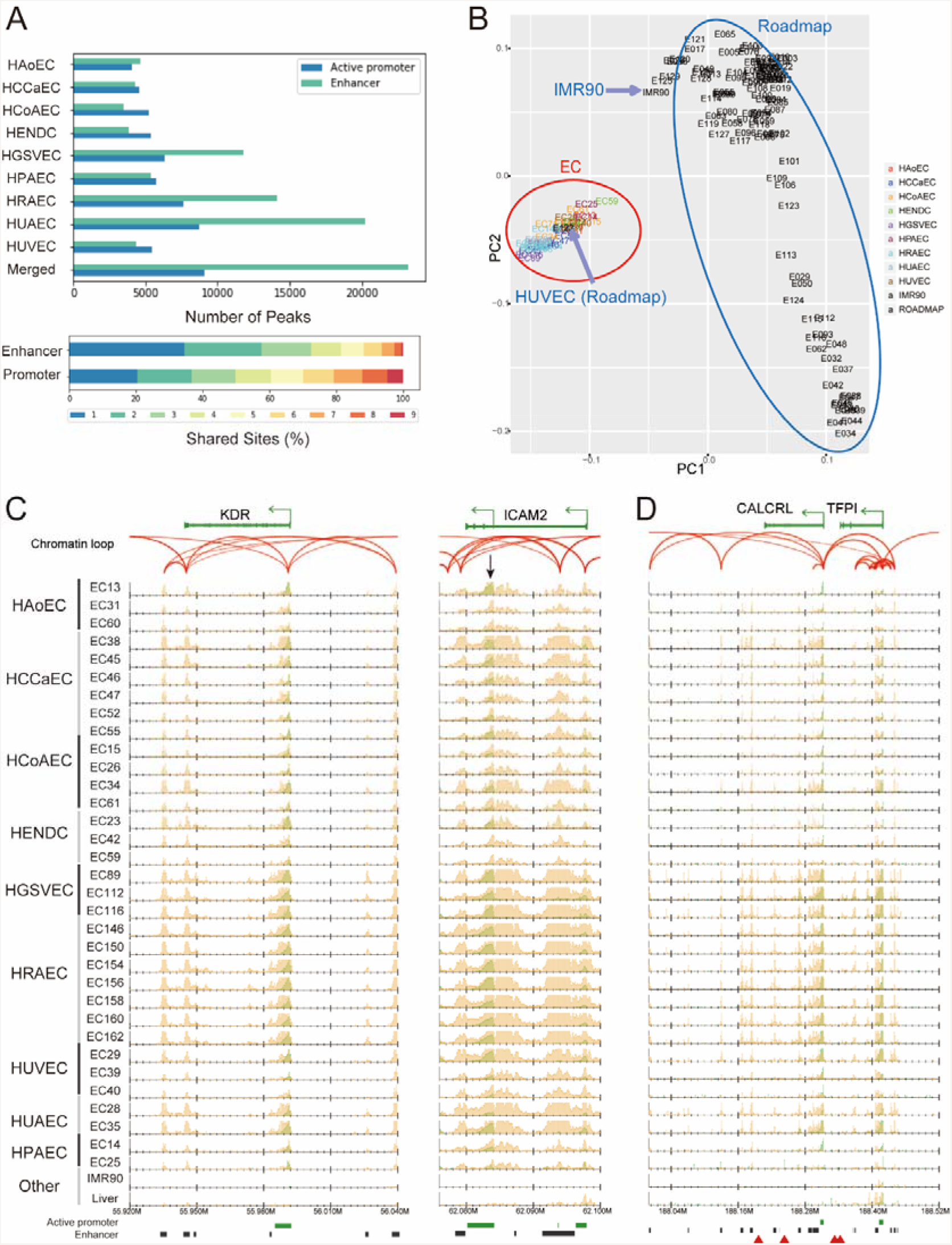
ChIP-seq data indicate variation in the chromatin status of ECs. (A) Top: The number of active promoter and enhancer sites for the nine cell types along with the merged reference sites. Bottom: The percentage of the reference active promoter and enhancer sites shared by one to nine of the EC types. (B) PCA plot using H3K27ac read densities. All EC samples in this paper (red circle) as well as 117 cell lines from Roadmap Epigenomics Project (blue circle) are shown. The label colors indicate the EC types. (C, D) Normalized read distribution of H3K4me3 (green) and H3K27ac (orange) in representative gene loci (C) *KDR* and *ICAM2* and (D) *CALCRL* and *TFPI* for all ECs and two other tissues (liver from the Roadmap and the IMR90 cells in this study). Chromatin loops based on ChIA-PET (read-pairs) are represented by red arches. Green bars, black bars and red triangles below each graph indicate active promoter sites, enhancer sites and GWAS SNPs, respectively.

### Evaluation of enhancer sites by PCA

To investigate the diverse distribution of our reference enhancer sites, we used the principal component analysis (PCA) based on the H3K27ac read densities in the integrated EC enhancer sites with the 117 cell lines from the Roadmap Epigenomics Project (14). We found that ECs were well clustered and separated from other cell lines (Figure 2B). Remarkably, HUVECs represented in the Roadmap Epigenomics Project dataset, termed E122, were properly included in the EC cluster (red circle). In contrast, IMR90 cells from our study were included in the non-EC cluster (blue circle). This result supported the reliability of our EC-specific enhancer profiling. It should be noted, however, that the samples for each EC cell type (indicated by different colors) were not well clustered, possibly because the EC type-specific difference is minuscule and is disrupted by differences at the level of the individual.

### Identification of enhancer-promoter interactions by ChIA-PET

We sought to identify the corresponding gene for the reference enhancer sites and used chromatin loop data obtained from the Chromatin Interaction Analysis by Paired-End Tag Sequencing (ChIA-PET) data using RNA Polymerase II (Pol II) in HUVECs. We identified 292 significant chromatin loops (false discovery rate (FDR) < 0.05), 49.3% (144 loops) of which connected promoter and enhancer sites. Even when we used all chromatin loops (at least one read pair), 27.4% (8,782 of 31,997) of them linked to enhancer-promoter sites. Remarkably, 48.1% (4,228 of 8,782) of loops connected the distal enhancer sites. In total, we identified 2,686 distal enhancer sites that are connected by chromatin loops. We also detected enhancer-enhancer (3,136, 9.8%) and promoter-promoter (11,618, 36.3%) loops, suggesting physically aggregated chromatin hubs in which multiple promoters and enhancers interact (15). As the ChIA-PET data are derived from RNA Pol II-associated loops in HUVECs, chromatin interactions in active genes could be detected.

### Identification of EC-specific sites

Next, we identified EC-specific enhancer sites by excluding any sites from our reference sites that overlapped with those of our IMR90 cells and other cell types from the Roadmap Epigenomics Project, except HUVECs (E122). As a result, we obtained 3,765 EC-specific enhancer sites (Supplementary Table S3), some of which were located around known marker genes of ECs with chromatin loops. One example is kinase insert domain receptor (*KDR*; Figure 2C, left), which functions as the VEGF receptor, causing endothelial proliferation, survival, migration, tubular morphogenesis and sprouting (16). The TSS of *KDR* was marked as an active promoter (enriched for both H3K4me3 and H3K27ac) and physically interacted with the EC-specific enhancer sites indicated by H3K27ac, ~50 kbp upstream and downstream of the TSS. Another example is intercellular adhesion molecule 2 (*ICAM2*, Figure 2C, right), which is an endothelial marker and is involved in the binding to white blood cells that occurs during the antigen-specific immune response (17). This gene has two known TSSs, both of which were annotated as active promoters in ECs, and one TSS that was EC specific (black arrow). This EC-specific TSS did not have a ChIA-PET interaction, and, likewise, the enhancer sites within the entire gene body did not directly interact with the adjacent promoter sites, implying the distinctive regulation of the two *ICAM2* promoters.

### Genome-wide association study (GWAS) enrichment analysis

To explore the correlation of EC-specific reference enhancer sites with sequence variants associated with disease phenotypes, we obtained reference GWAS single-nucleotide polymorphisms (SNPs) from the GWAS catalog (18) and identified significantly enriched loci by permutation analysis (19). Notably, we identified 67 enhancer sites that markedly overlapped with GWAS SNPs associated with “heart”, “coronary” and “cardiac” (Z score > 5.0, Supplementary Table S4). The most notable region was around *CALCRL* and *TFPI* loci (chr2:188146468-188248446, Figure 2D). The EC-specific enhancer region in these loci contained four GWAS risk variants (Figure 2D, red triangles), three of which are associated with coronary artery/heart disease (20, 21). Another example is the *RSPO3* locus (Supplementary Figure S4). The upstream distal enhancer regions of that gene contained four GWAS SNPs that are associated with cardiovascular disease and blood pressure (22, 23).

### Functional analysis of the reference sites

We next investigated whether any characteristic sequence feature is observed in the EC-specific enhancer and promoter sites (Supplementary Figure S5). We found a putative motif for EC-specific enhancer sites, which is closely similar to the canonical motifs of the homeobox genes *bcd*, *oc*, *Gsc* and *PITX1,2,3* (Figure 3). In fact, most of the EC-specific enhancer sites consisted of enhancers of HGSVECs (47.0%), HRAECs (37.4%) and HUAECs (68.3%) (Supplementary Figure S6), suggesting that these enhancers contain this motif.

**Figure 3.**
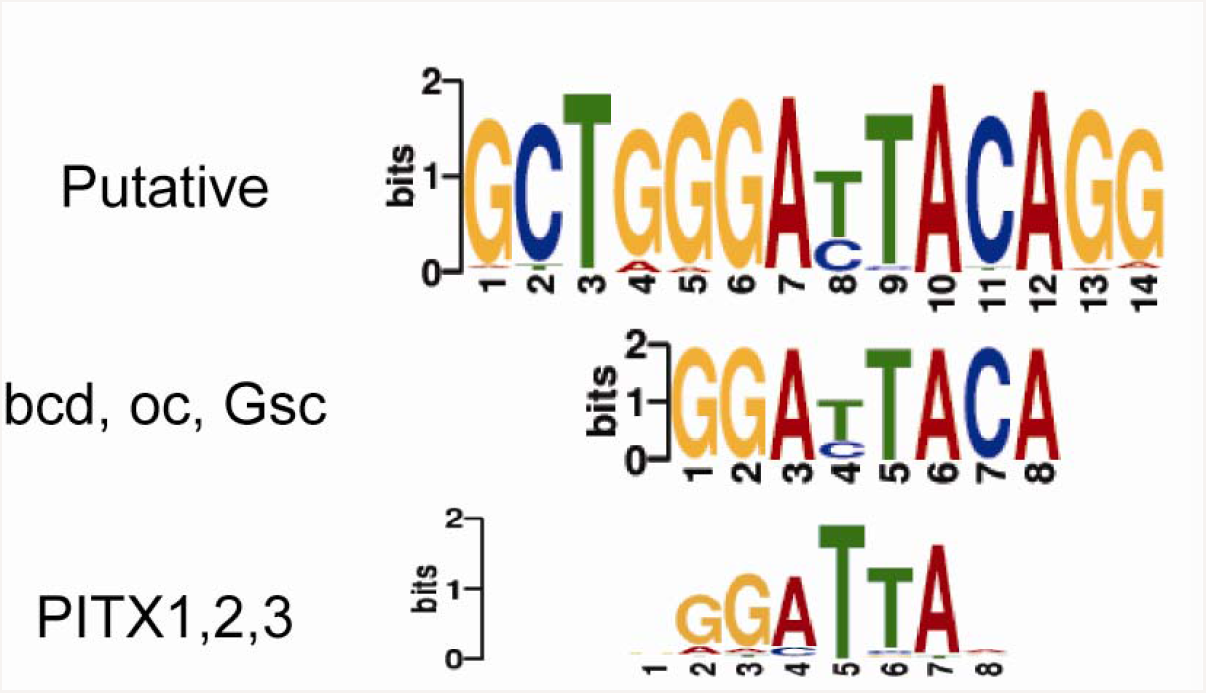
The identified *de novo* motif from EC-specific enhancer sites. The two related canonical motifs derived from JASPAR database are also shown.

We also looked into the Gene Ontology (GO) classifications under Biological Process for the enhancer sites using GREAT (24) and found that the enhancer sites (both all sites and EC-specific sites) have GO terms that are more specific to the vascular system (e.g., platelet activation, myeloid leukocyte activation and vasculogenesis), as compared with active promoter sites (e.g., mRNA catabolic process, Supplementary Figure S7). This also suggests that the enhancer sites are more likely to be associated with EC-specific functions, whereas promoter sites are also correlated with the more common biological functions.

### Differential analysis and clustering across EC types

One important issue of this study is to clarify the epigenomic/transcriptomic diversity across EC cell types. To circumvent variances at the level of the individual in each cell type observed (Figure 2C) and different S/N ratios, we fitted the value of peak intensity on the reference enhancer sites among samples using generalized linear models with the quantile normalization. By implementing a PCA, we confirmed that different cell samples in the same EC type were properly clustered (Figure 4A). The PCA also showed that different EC types can be divided into two subgroups based on the epigenomic landscape, corresponding to upper body (HAoEC, HCoAEC, HPAEC, HCCaEC and HENDC, purple circle) and lower body (HUVEC, HUAEC, HGSVEC and HRAEC) origins. A PCA based on gene expression data showed similar results to that based on the H3K27ac profile, although in the gene expression analysis HUAECs were more similar to heart ECs (Supplementary Figure S8).

**Figure 4:**
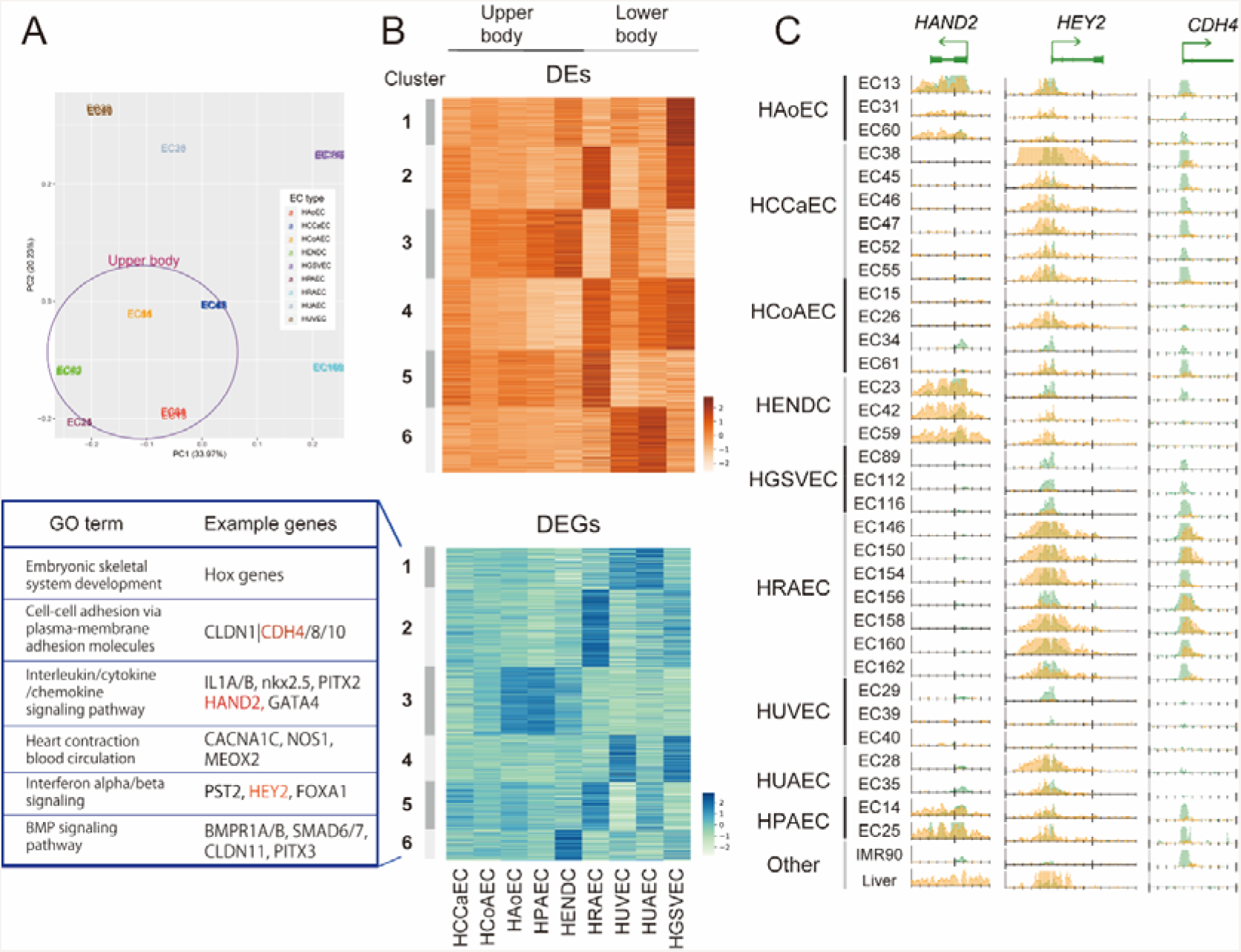
Comparative analysis of enhancer sites and gene expression across EC types. (A) PCA plot of EC samples based on H3K27ac read density fitted by generalized linear models. The color of samples indicates EC types. Samples from upper body are circled. (B) A k-means clustering (k = 6) analysis of DEs (upper) and DEGs (lower) across EC types (a representative for each type) based on Z-scores. The example genes and related GO terms obtained by Metascape (62) for DEG clusters are also shown. (C) Read distribution of H3K4me3 (green) and H3K27ac (orange) for the genes highlighted in red in panel (B).

To further investigate this tendency, we implemented a multiple-group differential analysis with respect to H3K4me3, H3K27ac and gene expression data to obtain sites and genes whose values were significantly varied between any of the nine cell types. With the threshold FDR < 1e-5, we identified 753 differential H3K4me3 sites (differential promoters, DPs; 8.3% from 9,121 active promoter sites), 2,979 differential H3K27ac sites (differential enhancers, DEs; 9.2% from 32,323 active promoter and enhancer sites) and 879 differentially expressed genes (DEGs; 2.1% from 41,880 genes). As expected, DPs and DEs were more enriched around DEGs, as compared with all genes. DPs were enriched within ~10 kbp from TSSs, whereas DEs were more broadly distributed (~100 kbp) (Supplementary Figure S9), indicative of the longer-range interactions between enhancers and their corresponding genes.

We then implemented k-means clustering (k = 6) to characterize the overall variability of DEGs, DEs (Figure 4B) and DPs (Supplementary Figure S10). Although k = 6 was empirically defined and might not be biologically optimal to classify the nine EC types, the results could capture the differential patterns. The upregulated genes were roughly categorized into upper and lower body–specific EC types (Figure 4B), even though diverse expression patterns were observed overall. In particular, the expression patterns of the EC types around the heart (HCoAEC, HAoEC and HPAEC) were similar (cluster 3 of DP and DEG), consistent with the anatomical proximity of these ECs. HENDCs had uniquely expressed genes (cluster 6 of DEGs). Considering that most of the DEGs and Des are cooperatively enriched in more than one EC type, these nine EC types may use distinct combinations of multiple genes, rather than exclusively expressed individual genes, for their specific phenotype.

### DEGs that contribute to EC functions

Our clustering analysis also identified important genes for EC functions as DEGs (Figure 4B). For example, heart and neural crest derivatives expressed 2 (*HAND2*) and GATA binding protein 4 (*GATA4*) were expressed in HAoECs, HENDCs and HPAECs (cluster 3). HAND2 physically interacts with GATA4 and the histone acetyltransferase p300 to form the enhanceosome complex, which regulates tissue-specific gene expression in the heart (25). Another example is hes related family bHLH transcription factor with YRPW motif 2 (*HEY2*, also called *Hrt2*), a positive marker for arterial EC specification (26), which was grouped to cluster 5 and was expressed specifically in aorta-derived ECs but not in vein-derived ECs (HUVECs and HGSVECs). HRAECs showed uniquely upregulated genes, including cadherin 4 (*CDH4*); the protein product of this gene mediates cell-cell adhesion, and mutation of this gene is significantly associated with chronic kidney disease in the Japanese population (27). Interestingly, at TSSs of *CDH4* and *HEY2* loci, H3K4me3 was also enriched in some EC types in which the genes were not expressed, whereas the H3K27ac enrichment pattern at TSSs was correlated with the expression level (Figure 4C). This variation in H3K4me3 with/without H3K27ac enrichment may reflect the competence of expression, which cannot be fully captured by gene expression analysis.

DEGs also contained several notable gene families. One example is the claudin family, a group of transmembrane proteins involved in barrier and pore formation (28). Whereas *CLDN5* has been reported as a major constituent of the brain EC tight junctions that make up the blood-brain barrier (29), we found that seven other genes belonging to the claudin family (*CLDN1*, *7, 10, 11, 12, 14* and *15*) were expressed in ECs, and their expression pattern varied across EC types (Supplementary Figure S11). For example, in HUVECs, *CLDN11* was highly expressed but *CLDN14* was not, although the two claudins share a similar function for cation permeability (30). These observations suggest that distinct usages of specific claudin proteins may result in different phenotypes with respect to vascular barrier function. Consequently, these DEGs are thus usable as a reference marker set for each EC type.

### Homeobox genes are highly differentially expressed across EC types

We also found that DEGs identified in our analysis contained genes that were not previously acknowledged as relevant to the different EC types. Most strikingly, quite a few homeobox (*HOX*) genes were differentially expressed (cluster 1 in Figure 4B and Figure 5A). The human genome has four *HOX* clusters (*HOXA*, *B*, *C* and *D*), each of which contains 9–11 genes essential for determining the body axes during embryonic development, as well as regulating cell proliferation and migration in diverse organisms (31). These genes are transcribed sequentially in both time and space, following their positions within each cluster (32). Figure 5A shows that genes in *HOX* clusters *A*, *B and D* were highly expressed in all EC types, except HENDCs, possibly because HENDCs are derived from cardiac neural crest, whereas the other EC types are derived from mesoderm (33). *HOXC* genes were moderately expressed in HRAECs, HUVECs and HUAECs, but not in the upper-body ECs. More interestingly, perhaps, *HOXD* genes were not expressed in HPAECs, despite their similar expression pattern relative to other EC types around the heart (Figure 4B). This result implies the distinct use of *HOX* paralogs, especially *HOXD* genes, in ECs. We also found that the 5’ *HOX* genes (blue bars in Figure 5A) tended to be selectively expressed in EC types derived from the lower body (HGSVECs, HRAECs, HUAECs and HUVECs). Considering the collinearity of their activation during axial morphogenesis, it is conceivable that the type-specific expression of *HOX* clusters, especially in 5’ *HOX* genes, reflects the developmental origin of EC types and that distinct activation of *HOX* genes collectively maintains the diversity of the circulatory system.

**Figure 5:**
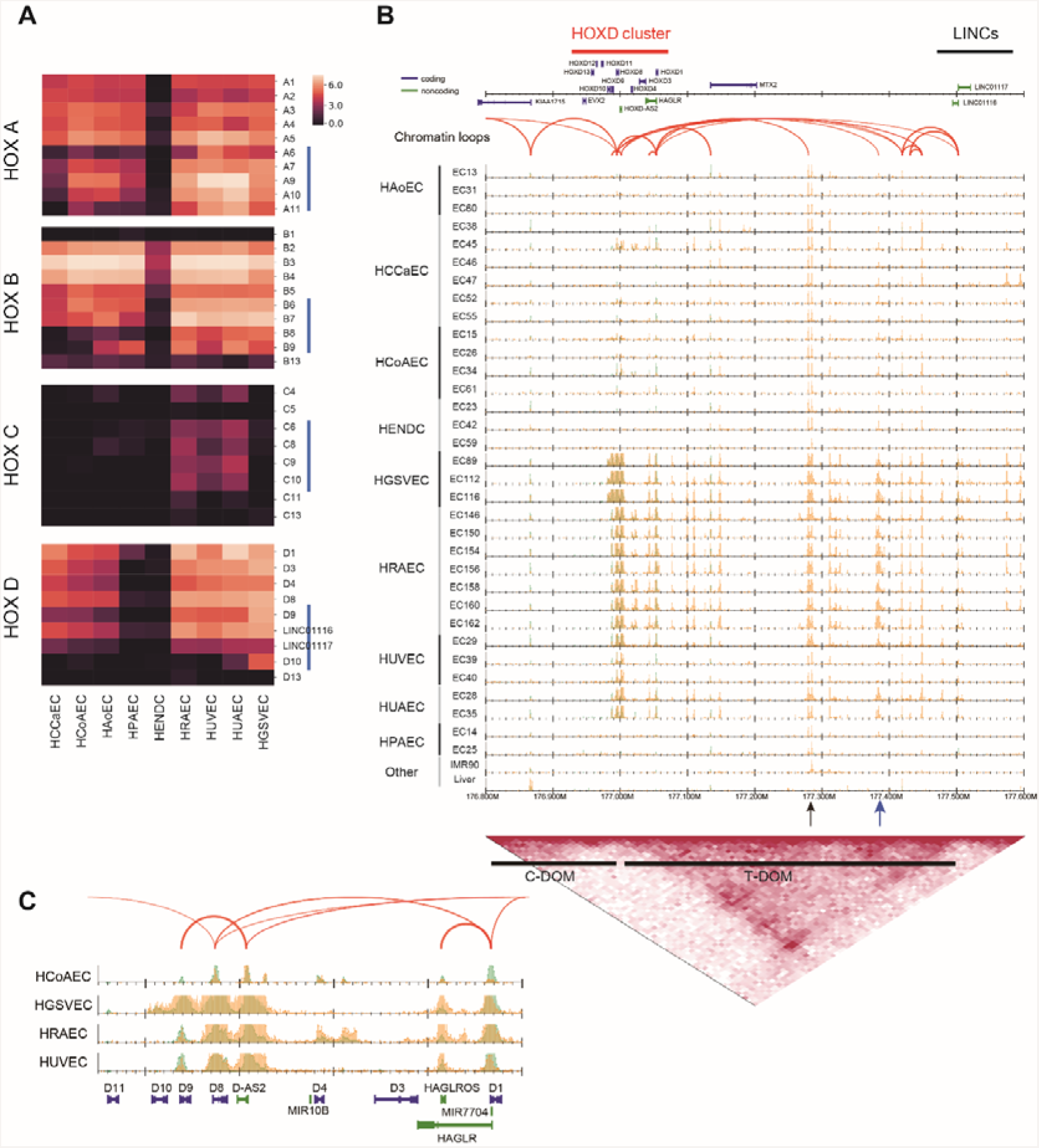
Differential expression of *HOX* genes. (A) Heatmaps visualizing the gene expression level (logged Transcripts Per Million (TPM)) of four *HOX* clusters and two long non-coding RNAs, LINC01117 (Hotdog) and LINC01116 (Twin of Hotdog). Blue vertical bars indicate the 5’ *HOX* genes. (B) Read distribution around the *HOXD* cluster (chr2: 176.8–177.6 Mbp). Bottom: topological interaction frequency, telomeric domain (T-DOM) and centromeric domain (C-DOM) identified by Hi-C data for HUVECs. (C) Comparison of read profiles around the *HOXD* region for four EC types.

It has been suggested that the more 3’ *HOX* genes tend to promote the angiogenic phenotype in ECs, whereas the more 5’ *HOX* genes tend to be inhibitory with respect to that phenotype (31). For example, *HOXD3* may promote wound healing and invasive or migratory behavior during angiogenesis in ECs (34). In contrast, *HOXD10* may function to inhibit EC migration by muting the downstream effects of other pro-angiogenic *HOX* genes (e.g., *HOX3* paralogs), and thus human ECs that overexpress *HOXD10* fail to form new blood vessels (35). Figure 5A shows that *HOXD10* was highly expressed in HGSVECs, which evokes the inhibition of the angiogenic phenotype as regulated by *HOXD10* in this cell type.

In addition to *HOX* genes, multiple non-*HOX* homeobox genes were also differentially regulated across ECs. For example, cluster 3 in the DEGs contained NK2 homeobox 5 (*Nkx-2.5*), which is essential for maintenance of ventricular identity (36); paired like homeodomain 2 (*PITX2*) and paired related homeobox 1 (*PRRX1*), which are both associated with the atrial fibrillation and cardioembolic ischemic stroke variants loci (37–39); and Meis homeobox 1 (*MEIS1*), which is required for heart development in mice (40). These were all associated with the GO term “blood vessel morphogenesis (GO:0048514)”. Interestingly, *PITX3* was mainly expressed in HENDCs (cluster 6), unlike *PITX2*. Another example is Mesenchyme Homeobox 2 (*MEOX2*, also known as Gax; cluster 4), which regulates senescence and proliferation in HUVECs (41) and was also expressed in HUAECs and HGSVECs but not in other cell types. Taken together with the finding that some binding motifs of homeobox genes including *PITX* were identified among the EC-specific enhancer sites (Figure 3), these data suggest that distinct combinations of proteins coded by *HOX* and non-*HOX* homeobox genes play a key role in mature human ECs for angiogenesis, vasculogenesis and wound healing, in addition to their function during the development and proliferation of ECs.

### Enhancers in the telomeric domain were upregulated within the *HOXD* cluster

Lastly, we investigated the epigenomic landscape around the *HOXD* cluster (Figure 5B). It has been reported that the mammalian *HOXD* cluster is located between two enhancer-rich topologically associating domains (TADs), the centromeric domain (C-DOM) and the telomeric domain (T-DOM), which are activated during limb and digit development, respectively (42). By using public Hi-C (genome-wide Chromosome Conformation Capture (3C)) data for HUVECs (43) to detect the T-DOM and C-DOM (bottom black bars), we observed the presence of EC enhancers in the T-DOM (Figure 5B), as in early stages of limb development (42). Of note, two long non-coding RNAs, *LINC01117* (*Hotdog*) and *LINC01116* (*Twin of Hotdog*), which physically contact the expressed *HOXD* genes and are activated during cecum budding (44), had ChIA-PET loops and showed similar expression patterns with *HOXD* genes in ECs (Figure 5A and 5B). In the T-DOM, some enhancers are likely to be activated in most EC types (Figure 5B, black arrow), whereas others are active only in ECs from the lower body (blue arrow), suggesting a physical interaction between these enhancers and each *HOXD* gene in a constitutive and a cell type–specific manner, respectively. A detailed view of the genomic region from *HOXD1* to *HOXD11* (Figure 5C) shows that H3K4me3 and H3K27ac are specifically enriched within the *HOXD10* locus in HGSVECs, which is consistent with their gene expression pattern. Because the *HOXD10* locus did not have ChIA-PET loops in HUVECs, there might be HGSVEC-specific chromatin loops.

## DISCUSSION

In this study, we analyzed the epigenomic status of the active histone modifications H3K4me3 and H3K27ac in 33 samples from nine different EC types by ChIP-seq, RNA-seq, ChIA-PET and Hi-C analyses. The integrative ChIP-seq analysis based on the samples from the human tissues of multiple donors is hampered by both individual variations and technical noise derived from tissue sample acquisition under various conditions (e.g., race, sex, age and the sample acquirement process), compared with the smaller difference among EC types. To overcome these issues, at least in part, we developed a robust procedure for comparative epigenome analysis, combined with chromatin interaction data. We successfully identified 3,765 EC-specific enhancer sites, 67 of which were highly significantly overlapping with GWAS SNPs. We aim to expand this analysis to other core histone marks including suppressive markers (e.g., H3K27me3 and H3K9me3) and apply semi-automated genome annotation methods (45). Because this type of genome annotation strategy with its associated assembling of broad marks is more sensitive to noise, more stringent quality control of tissue data will be required.

The PCA showed that EC types can be divided into those from the upper and from the lower body (Figure 4A). The nine EC types tend to use distinct combinations of multiple genes, rather than exclusively expressed genes, for their specific phenotype (Figure 4B and Figure 5A). Our results identified key marker genes that were differentially expressed across EC types, such as homeobox genes. The importance of several *HOX* genes for early vascular development and adult angiogenesis in pathological conditions has been reported (46). However, a systematic analysis of the regulation and roles of homeobox genes in mature tissue cells has been lacking. We identified a regulatory motif enriched in EC-specific enhancers that is very similar to those of *homeobox* genes and distinct epigenome states and chromatin conformations of *HOX* gene clusters and flanking regions in different EC types. Taken together, our data suggest the distinct roles and combinatorial usage of *HOX* genes during development and in regulating EC phenotypes throughout the body.

## CONCLUSION

The primary goal of the IHEC project is to generate high-quality reference epigenomes and make them available to the scientific community (10). To this end, we established an epigenetic catalog of various human ECs and implemented comprehensive analysis to elucidate the diversity of the epigenomic and transcriptomic landscape across EC types. The dataset presented in this study will be an important resource for future work on understanding the human cardiovascular system and its associated diseases.

## METHODS

### Tissue preparation

ECs were isolated from the vasculature and maintained as primary cultures, as reported (47, 48). Briefly, HAoECs, HCoAECs, HENDCs, HPAECs and HUVECs were isolated from the various vessels by incubating the vessels with collagenase at 37°C for 30 minutes. The aortic root was used for HAoEC isolation. Cells were plated in tissue culture-treated flasks (Iwaki Glass Co. Ltd., cat. No.) and cultured for one or more passages in modified VascuLife VEGF Endotnelial Medium (Lifeline Cell Technology). A reduced concentration of VEGF that was lower than 5 ng/mL was tested in preliminary cell culture experiments and then optimized to be as low as possible considering cell growth and viability (data not shown). The VEGF concentration was lowered to 250 pg/mL, which is lower than the standard culture conditions, to more closely replicate in vivo concentrations (49). ECs were separated from non-ECs using immunomagnetic beads. Fibroblasts were first removed using anti-fibroblast beads and the appropriate magnetic column (Miltenyi Biotec). The remaining cells were then purified using Dynabeads and anti-CD31 (BAM3567, R&D Systems). When positive selection was used, the bead-bound cells were removed from the cell suspension prior to cryopreservation.

HCCaECs and HRECs were prepared by an explant culture method (48). HGSVECs were isolated from discarded veins taken from patients at Saitama Medical University International Medical Center.

Quality control was performed using a sterility test (for bacteria, yeast and fungi), a PCR-based sterility test (for hepatitis B and C, HIV-I and -II and mycoplasma) and immunostaining-based characterization for von Willebrand factor (vWF) (>95% cells are positively stained (50)) and alpha-actin, and viability was determined by both counting and trypan blue staining.

### Cell culture

Purified ECs were cultured in VascuLife VEGF Endothelial Medium (Lifeline Cell Technology) with 250 pg/mL VEGF. Cells were maintained at 37°C in a humidified 5% CO_2_ incubator, and the medium was changed every 3 days. The cells used in the experiments were from passage 6 or less. The cryopreservation solution used consisted of VascuLife VEGF Endothelial medium, containing 250 pg/mL VEGF, 12% Fetal Bovine Serum (FBS) and 10% Dimethylsulfoxide (DMSO).

### RNA-seq analysis

Poly(A)-containing mRNA molecules were isolated from total RNA and then converted to cDNA with oligo(dT) primers using a TruSeq RNA Sample Preparation kit v2 (Illumina) and were sequenced with a HiSeq 2500 system (Illumina). We applied sequenced paired-end reads to kallisto version 0.43.1 (51) with the “--rf-stranded -b 100” option, which estimates the transcript-level expression values as Transcripts Per Kilobase Million (TPM, Ensembl gene annotation GRCh37). These transcript-level expression values were then assembled to the gene-level by tximport (52). We also obtained RNA-seq data from IMR90 cells from the Sequence Read Archive (SRA) (www.ncbi.nlm.nih.gov/sra) under accession number SRR2952390. The full list of gene expression data is available at the NCBI Gene Expression Omnibus (GEO) under the accession number GSE131953.

### ChIP

For each EC sample, two million ECs were plated on a 15-cm culture plate and cultured until confluency. The cells were crosslinked for 10 minutes using 1% paraformaldehyde. After quenching using 0.2 M glycine, cells were collected using a scraper, resuspended in SDS lysis buffer (10 mM Tris-HCl, 150 mM NaCl, 1% SDS, 1 mM EDTA; pH 8.0) and fragmented by sonication (Branson, Danbury, CT, USA; 10 minutes). Samples were stored at –80°C before use. To perform ChIP, antibodies against histone modifications (CMA304 and CMA309 for H3K4me3 and H3K27ac, respectively) (53) were used in combination with protein G Sepharose beads (GE Healthcare Bio-Sciences AB, Sweden). The prepared DNA was quantified using Qubit (Life Technologies/Thermo Fisher Scientific), and >10 ng of DNA was processed, as described below. The primer sequences for ChIP-qPCR were as follows: for H3K4me3, KDR (Fw: CCACAGACTCGCTGGGTAAT, Rv: GAGCTGGAGAGTTGGACAGG) and GAPDH (Fw: CGCTCACTGTTCTCTCCCTC, Rv: GACTCCGACCTTCACCTT CC); for H3K27ac, ANGPTL4 (Fw: TAGGGGAATGGGTAGGGAAG, Rv: AGTTCTCAGGCAGGTGGAGA) and GATA2 (Fw: AGACGACCCCAACTGACATC, Rv: CCTTCAAATGCAGACGCTTT) and, as a negative control, HBB (Fw: GGGCTGAGGGTTTGAAGTCC, Rv: CATGGTGTCTGTTTGAGGTTGC).

### ChIP-seq analysis

Sequencing libraries were made using the NEBNext ChIP-Seq Library Prep Master Mix Set of Illumina (New England Biolabs). Sequenced reads were mapped to the human genome using Bowtie version 1.2 (54) allowing two mismatches in the first 28 bases per read and outputting only uniquely mapped reads (−n2 −m1 option). Peaks were called by DROMPA version 3.5.1 (55) using the stringent parameter set (-sm 200-pthre_internal 0.00001 -pthre_enrich 0.00001) to mitigate the effect of technical noise. The mapping and peak statistics are summarized in Supplementary Table S2.

### Quality validation of ChIP-seq samples

We checked the quality of each sample based on the peak number, library complexity and GC content bias by DROMPA; the normalized strand coefficient and background uniformity by SSP (12); inter-sample correlation (jaccard index of peak overlap) by bedtools (https://github.com/arq5x/bedtools2); and the pairwise correlations of read coverage by deepTools version 2.5.0 (56) (Supplementary Figure S1).

### Regression analysis of ChIP-seq data

To estimate the expression level of a gene from the level of its histone modifications, we implemented the linear regression analysis proposed by Karlic *et al.* (13) with minor modifications. We built a two-variable model to predict the expression level for each mRNA as follows:

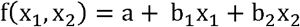

where x_1_ and x_2_ are the log-scale basepair coverage in a region of 4-kbp surrounding the TSSs covered by obtained peaks of H3K4me3 and H3K27ac, respectively. We used the level of protein-coding mRNA in autosomes as an estimation of the level of histone modifications. We then learned the parameters *a*, *b*_*1*_ and *b*_*2*_ using all of the EC samples and the IMR90 sample to minimize the differences between observed and expected values. Using the learned parameter set, we predicted the expression value for each mRNA and calculated the Pearson correlation between observed and expected values.

### Definition of reference promoter and enhancer sites

As shown in Figure 1B, we defined active promoters and enhancers as “H3K4me3 sites overlapping with H3K27ac sites by ≥1 bp” and “H3K27ac sites not overlapping with H3K4me3 sites”, respectively, based on the annotation of the Roadmap Epigenomics consortium (14). Peaks from sex chromosomes were excluded to ignore sex-specific difference. To avoid the effect of individual differences, the common sites among all samples were used as the reference sites for each cell type. Then the reference sites of all cell types were merged into the reference sites of ECs. Multiple sites that were within 100 bp of each other were merged to avoid multiple counts of large individual sites. The generated reference promoter and enhancer sites are available at the GEO under the accession number GSE131953.

### Identification of EC-specific sites

We called peaks for H3K4me3 and H3K27ac for all 117 cell lines in the Roadmap Epigenomics Project by DROMPA with the same parameter set and excluded the sites in the reference promoter and enhancer sites of ECs that overlapped the H3K4me3 peaks (promoter sites) or H3K27ac peaks (enhancer sites) of all cells except for E122 (HUVECs) from the Roadmap Epigenomics Project. Similarly, we further excluded the sites that overlapped H3K27ac peaks of IMR90 cells generated by this study, to avoid the protocol-dependent false-positive peaks. The resulting sites were used as EC-specific sites. We also defined “distal enhancer sites” as those that are >10 kbp from the nearest TSS. These sites are summarized in Table S2.

### GWAS enrichment analysis

We implemented GWAS enrichment analysis using a strategy similar to that of Lake *et al.* (19). We obtained reference SNPs from the GWAS Catalog [18]. We then calculated the occurrence probability of GWAS SNPs associated with the terms “heart”, “coronary” and “cardiac” in 100-kb regions centered on all EC-specific enhancer sites and investigated their statistical significance by random permutations. We extended each enhancer site to a 100-kb region to consider linkage disequilibrium with GWAS SNPs. We identified the enhancer sites with a Z-score > 5.0. We shuffled the enhancer sites randomly within each chromosome, ignoring the centromeric region, using bedtools shuffle command.

### Differential analysis of multiple groups for histone modification and gene expression

We applied the ANOVA-like test in edgeR (57) based on the normalized read counts of H3K4me3 ChIP-seq data in active promoters and H3K27ac ChIP-seq data in active promoters and enhancers, as well as gene expression data, while fitting the values among samples to estimate dispersion using generalized linear models. For RNA-seq data, the count data were fitted using a generalized linear model, and the Z-score was calculated based on logged values. For ChIP-seq data, we also applied the quantile normalization to peak intensity in advance of the fitting because this model does not consider the different S/N ratios among samples (58). This normalization assumes that the S/N ratio for most of the common peaks should be the same among all samples in which the same antibody was used. Supplementary Figure S12 shows the distribution patterns of the H3K27ac read density normalized for quantile normalization for all ECs.

### Chromatin interaction analysis

We used ChIA-PET data mediated by RNA Pol II for HUVECs (59). We acquired fastq files from the GEO under accession number GSE41553, applied Mango (60) with default parameter settings and identified the 943 significant interactions (1,886 sites, FDR < 0.05). For Hi-C analysis, we acquired .hic files for HUVECs from the GEO under accession number GSE63525 and applied Juicer (43) to obtained the TAD structure (Figure 5B).

### Motif analysis

We used MEME-ChIP version 5.0.1 (61) with the parameter set “-meme-mod zoops -meme-minw 6-meme-maxw 14” with the motif data “JASPAR2018_CORE_non-redundant.meme”.

## List of abbreviations

EC: Endothelial cells
S/N: signal-to-noise ratio
PCA: principal component analysis
Pol II: RNA Polymerase II
FDR: false discovery rate
GWAS: Genome-wide association study
SNP: single-nucleotide polymorphism
GO: Gene Ontology
DP: differential promoter
DE: differential enhancer
DEG: differentially expressed gene
TAD: topologically associating domain
C-DOM: the centromeric domain
T-DOM: the telomeric domain
vWF: von Willebrand factor
TPM: Transcripts Per Kilobase Million

## Declarations

### Ethics approval and consent to participate

Human clinical specimens were prepared from discarded tissue during surgery under consent of donors at Saitama Medical University International Medical Center according to the Institutional Review Board (IRB) protocol 15-209, at Ohta Memorial Hospital according to the IRB protocols 068 and 069 and at The University of Tokyo according to the IRB protocol G3577. Primary ECs were isolated at The University of Tokyo according to the IRB protocol 17-311. Other ECs were prepared from a commercial biobank (Lifeline Cell Technology, Frederick, MD). DNA and RNA samples were prepared at The University of Tokyo according to the IRB protocol 12-81.

### Consent for publication

As to purchased cells, not applicable. As to primary cultivated cells from human tissue, written informed consent was obtained from the patients for publication of their individual details and accompanying images in this manuscript. The consent form is held by the authors and is available for review by the Editor-in-Chief.

### Availability of data and materials

The raw sequencing data and processed files are available at the GEO under the accession numbers GSE131953 (ChIP-seq) and GSE131681 (RNA-seq) with links to BioProject accession number PRJNA532996. The data summary, quality control results, the full list of ChIA-PET loops and visualization figures are available on the EC analysis website (https://rnakato.github.io/HumanEndothelialEpigenome/).

### Authors’ contributions

R. Nakato designed the studies, performed bioinformatics analysis and drafted the manuscript. Y.W. designed the studies, carried out the molecular genetic studies and drafted the manuscript. R. Nakaki, N.N., S.T., T. Kohro, N.O., Y.S. and T.M. performed bioinformatics analyses. G.N. and K.T. performed mRNA-seq. Y. Katou performed library preparation and sequencing. Y. Kanki performed cell culture, prepared samples and drafted the manuscript. M.K. and A.K. performed cell culture and prepared samples. Y.H-T. and A.I-T. prepared reagents and performed chromatin immunoprecipitation. H.F., A.I., H.N., M.N., T.S., S.N. H.W., S.O., M.A., R.C.M., K.W.H., T. Kawakatsu, M.G., H.Y., H. Kume, and Y.H. prepared primary cell cultures. H.A. and K.S. designed the studies and drafted the manuscript. H.Kimura prepared antibodies, designed the studies and drafted the manuscript.

## Acknowledgements

Advanced Medical Graphics (MA, USA) prepared the graphics of the organs and tissues. Niinami Hiroshi continuously supported domestic sample collection and preparation. We thank Ryozo Omoto for providing the pipeline from the surgical operating room to the research laboratory.

## References

1. Aird WC. Phenotypic heterogeneity of the endothelium: II. Representative vascular beds. Circulation research. 2007;100(2):174–90.

2. Thompson RC, Allam AH, Lombardi GP, Wann LS, Sutherland ML, Sutherland JD, et al. Atherosclerosis across 4000 years of human history: the Horus study of four ancient populations. Lancet. 2013;381(9873):1211–22.

3. Eppihimer MJ, Wolitzky B, Anderson DC, Labow MA, Granger DN. Heterogeneity of expression of E- and P-selectins in vivo. Circulation research. 1996;79(3):560–9.

4. Kaufer E, Factor SM, Frame R, Brodman RF. Pathology of the radial and internal thoracic arteries used as coronary artery bypass grafts. The Annals of thoracic surgery. 1997;63(4):1118–22.

5. Aird WC. Phenotypic heterogeneity of the endothelium: I. Structure, function, and mechanisms. Circulation research. 2007;100(2):158–73.

6. Sabik JF, 3rd, Raza S, Blackstone EH, Houghtaling PL, Lytle BW. Value of internal thoracic artery grafting to the left anterior descending coronary artery at coronary reoperation. Journal of the American College of Cardiology. 2013;61(3):302–10.

7. Chi JT, Chang HY, Haraldsen G, Jahnsen FL, Troyanskaya OG, Chang DS, et al. Endothelial cell diversity revealed by global expression profiling. Proceedings of the National Academy of Sciences of the United States of America. 2003;100(19):10623–8.

8. Kanki Y, Kohro T, Jiang S, Tsutsumi S, Mimura I, Suehiro J, et al. Epigenetically coordinated GATA2 binding is necessary for endothelium-specific endomucin expression. The EMBO journal. 2011;30(13):2582–95.

9. Tozawa H, Kanki Y, Suehiro J, Tsutsumi S, Kohro T, Wada Y, et al. Genome-wide approaches reveal functional interleukin-4-inducible STAT6 binding to the vascular cell adhesion molecule 1 promoter. Mol Cell Biol. 2011;31(11):2196–209.

10. Stunnenberg HG, International Human Epigenome C, Hirst M. The International Human Epigenome Consortium: A Blueprint for Scientific Collaboration and Discovery. Cell. 2016;167(5):1145–9.

11. Ernst J, Kheradpour P, Mikkelsen TS, Shoresh N, Ward LD, Epstein CB, et al. Mapping and analysis of chromatin state dynamics in nine human cell types. Nature. 2011;473(7345):43–9.

12. Nakato R, Shirahige K. Sensitive and robust assessment of ChIP-seq read distribution using a strand-shift profile. Bioinformatics. 2018;34(14):2356–63.

13. Karlic R, Chung HR, Lasserre J, Vlahovicek K, Vingron M. Histone modification levels are predictive for gene expression. Proceedings of the National Academy of Sciences of the United States of America. 2010;107(7):2926–31.

14. Roadmap Epigenomics Consortium, Kundaje A, Meuleman W, Ernst J, Bilenky M, Yen A, et al. Integrative analysis of 111 reference human epigenomes. Nature. 2015;518(7539):317–30.

15. Allahyar A, Vermeulen C, Bouwman BAM, Krijger PHL, Verstegen M, Geeven G, et al. Enhancer hubs and loop collisions identified from single-allele topologies. Nature genetics. 2018;50(8):1151–60.

16. Waltenberger J, Claesson-Welsh L, Siegbahn A, Shibuya M, Heldin CH. Different signal transduction properties of KDR and Flt1, two receptors for vascular endothelial growth factor. The Journal of biological chemistry. 1994;269(43):26988–95.

17. Lyck R, Enzmann G. The physiological roles of ICAM-1 and ICAM-2 in neutrophil migration into tissues. Current opinion in hematology. 2015;22(1):53–9.

18. Buniello A, MacArthur JAL, Cerezo M, Harris LW, Hayhurst J, Malangone C, et al. The NHGRI-EBI GWAS Catalog of published genome-wide association studies, targeted arrays and summary statistics 2019. Nucleic Acids Res. 2019;47(D1):D1005–D12.

19. Lake BB, Chen S, Sos BC, Fan J, Kaeser GE, Yung YC, et al. Integrative single-cell analysis of transcriptional and epigenetic states in the human adult brain. Nature biotechnology. 2018;36(1):70–80.

20. van der Harst P, Verweij N. Identification of 64 Novel Genetic Loci Provides an Expanded View on the Genetic Architecture of Coronary Artery Disease. Circulation research. 2018;122(3):433–43.

21. Coronary Artery Disease Genetics C. A genome-wide association study in Europeans and South Asians identifies five new loci for coronary artery disease. Nature genetics. 2011;43(4):339–44.

22. Kichaev G, Bhatia G, Loh PR, Gazal S, Burch K, Freund MK, et al. Leveraging Polygenic Functional Enrichment to Improve GWAS Power. Am J Hum Genet. 2019;104(1):65–75.

23. Ehret GB, Ferreira T, Chasman DI, Jackson AU, Schmidt EM, Johnson T, et al. The genetics of blood pressure regulation and its target organs from association studies in 342,415 individuals. Nat Genet. 2016;48(10):1171–84.

24. McLean CY, Bristor D, Hiller M, Clarke SL, Schaar BT, Lowe CB, et al. GREAT improves functional interpretation of cis-regulatory regions. Nature biotechnology. 2010;28(5):495–501.

25. Dai YS, Cserjesi P, Markham BE, Molkentin JD. The transcription factors GATA4 and dHAND physically interact to synergistically activate cardiac gene expression through a p300-dependent mechanism. The Journal of biological chemistry. 2002;277(27):24390–8.

26. Aranguren XL, Agirre X, Beerens M, Coppiello G, Uriz M, Vandersmissen I, et al. Unraveling a novel transcription factor code determining the human arterial-specific endothelial cell signature. Blood. 2013;122(24):3982–92.

27. Yoshida T, Kato K, Yokoi K, Oguri M, Watanabe S, Metoki N, et al. Association of genetic variants with chronic kidney disease in Japanese individuals with or without hypertension or diabetes mellitus. Exp Ther Med. 2010;1(1):137–45.

28. Morita K, Furuse M, Fujimoto K, Tsukita S. Claudin multigene family encoding four-transmembrane domain protein components of tight junction strands. Proc Natl Acad Sci U S A. 1999;96(2):511–6.

29. Morita K, Sasaki H, Furuse M, Tsukita S. Endothelial claudin: claudin-5/TMVCF constitutes tight junction strands in endothelial cells. The Journal of cell biology. 1999;147(1):185–94.

30. Amasheh S, Fromm M, Gunzel D. Claudins of intestine and nephron - a correlation of molecular tight junction structure and barrier function. Acta physiologica. 2011;201(1):133–40.

31. Gorski DH, Walsh K. Control of vascular cell differentiation by homeobox transcription factors. Trends in cardiovascular medicine. 2003;13(6):213–20.

32. Kmita M, Duboule D. Organizing axes in time and space; 25 years of colinear tinkering. Science. 2003;301(5631):331–3.

33. Srivastava D. Making or breaking the heart: from lineage determination to morphogenesis. Cell. 2006;126(6):1037–48.

34. Uyeno LA, Newman-Keagle JA, Cheung I, Hunt TK, Young DM, Boudreau N. Hox D3 expression in normal and impaired wound healing. The Journal of surgical research. 2001;100(1):46–56.

35. Myers C, Charboneau A, Cheung I, Hanks D, Boudreau N. Sustained expression of homeobox D10 inhibits angiogenesis. The American journal of pathology. 2002;161(6):2099–109.

36. Targoff KL, Colombo S, George V, Schell T, Kim SH, Solnica-Krezel L, et al. Nkx genes are essential for maintenance of ventricular identity. Development. 2013;140(20):4203–13.

37. Christophersen IE, Rienstra M, Roselli C, Yin X, Geelhoed B, Barnard J, et al. Large-scale analyses of common and rare variants identify 12 new loci associated with atrial fibrillation. Nature genetics. 2017;49(6):946–52.

38. Gudbjartsson DF, Arnar DO, Helgadottir A, Gretarsdottir S, Holm H, Sigurdsson A, et al. Variants conferring risk of atrial fibrillation on chromosome 4q25. Nature. 2007;448(7151):353–7.

39. Neurology Working Group of the Cohorts for H, Aging Research in Genomic Epidemiology Consortium tSGN, the International Stroke Genetics C. Identification of additional risk loci for stroke and small vessel disease: a meta-analysis of genome-wide association studies. The Lancet Neurology. 2016;15(7):695–707.

40. Wang H, Liu C, Liu X, Wang M, Wu D, Gao J, et al. MEIS1 Regulates Hemogenic Endothelial Generation, Megakaryopoiesis, and Thrombopoiesis in Human Pluripotent Stem Cells by Targeting TAL1 and FLI1. Stem Cell Reports. 2018;10(2):447–60.

41. Gohn CR, Blue EK, Sheehan BM, Varberg KM, Haneline LS. Mesenchyme Homeobox 2 Enhances Migration of Endothelial Colony Forming Cells Exposed to Intrauterine Diabetes Mellitus. Journal of cellular physiology. 2017;232(7):1885–92.

42. Andrey G, Montavon T, Mascrez B, Gonzalez F, Noordermeer D, Leleu M, et al. A switch between topological domains underlies HoxD genes collinearity in mouse limbs. Science. 2013;340(6137):1234167.

43. Rao SS, Huntley MH, Durand NC, Stamenova EK, Bochkov ID, Robinson JT, et al. A 3D map of the human genome at kilobase resolution reveals principles of chromatin looping. Cell. 2014;159(7):1665–80.

44. Delpretti S, Montavon T, Leleu M, Joye E, Tzika A, Milinkovitch M, et al. Multiple enhancers regulate Hoxd genes and the Hotdog LncRNA during cecum budding. Cell reports. 2013;5(1):137–50.

45. Hoffman MM, Buske OJ, Wang J, Weng Z, Bilmes JA, Noble WS. Unsupervised pattern discovery in human chromatin structure through genomic segmentation. Nature methods. 2012;9(5):473–6.

46. Kachgal S, Mace KA, Boudreau NJ. The dual roles of homeobox genes in vascularization and wound healing. Cell adhesion & migration. 2012;6(6):457–70.

47. Wada Y, Sugiyama A, Yamamoto T, Naito M, Noguchi N, Yokoyama S, et al. Lipid accumulation in smooth muscle cells under LDL loading is independent of LDL receptor pathway and enhanced by hypoxic conditions. Arteriosclerosis, thrombosis, and vascular biology. 2002;22(10):1712–9.

48. Kobayashi M, Inoue K, Warabi E, Minami T, Kodama T. A simple method of isolating mouse aortic endothelial cells. Journal of atherosclerosis and thrombosis. 2005;12(3):138–42.

49. Vermeulen PB, Salven P, Benoy I, Gasparini G, Dirix LY. Blood platelets and serum VEGF in cancer patients. Br J Cancer. 1999;79(2):370–3.

50. Au-Yeung KK, Woo CW, Sung FL, Yip JC, Siow YL, O K. Hyperhomocysteinemia activates nuclear factor-kappaB in endothelial cells via oxidative stress. Circ Res. 2004;94(1):28–36.

51. Bray NL, Pimentel H, Melsted P, Pachter L. Near-optimal probabilistic RNA-seq quantification. Nature biotechnology. 2016;34(5):525–7.

52. Soneson C, Love MI, Robinson MD. Differential analyses for RNA-seq: transcript-level estimates improve gene-level inferences. F1000Research. 2015;4:1521.

53. Kimura H, Hayashi-Takanaka Y, Goto Y, Takizawa N, Nozaki N. The organization of histone H3 modifications as revealed by a panel of specific monoclonal antibodies. Cell structure and function. 2008;33(1):61–73.

54. Langmead B, Trapnell C, Pop M, Salzberg SL. Ultrafast and memory-efficient alignment of short DNA sequences to the human genome. Genome biology. 2009;10(3):R25.

55. Nakato R, Itoh T, Shirahige K. DROMPA: easy-to-handle peak calling and visualization software for the computational analysis and validation of ChIP-seq data. Genes to cells: devoted to molecular & cellular mechanisms. 2013;18(7):589–601.

56. Ramirez F, Ryan DP, Gruning B, Bhardwaj V, Kilpert F, Richter AS, et al. deepTools2: a next generation web server for deep-sequencing data analysis. Nucleic acids research. 2016;44(W1):W160–5.

57. Zhou X, Lindsay H, Robinson MD. Robustly detecting differential expression in RNA sequencing data using observation weights. Nucleic acids research. 2014;42(11):e91.

58. Nakato R, Shirahige K. Recent advances in ChIP-seq analysis: from quality management to whole-genome annotation. Briefings in bioinformatics. 2017;18(2):279–90.

59. Papantonis A, Kohro T, Baboo S, Larkin JD, Deng B, Short P, et al. TNFalpha signals through specialized factories where responsive coding and miRNA genes are transcribed. The EMBO journal. 2012;31(23):4404–14.

60. Phanstiel DH, Boyle AP, Heidari N, Snyder MP. Mango: a bias-correcting ChIA-PET analysis pipeline. Bioinformatics. 2015;31(19):3092–8.

61. Machanick P, Bailey TL. MEME-ChIP: motif analysis of large DNA datasets. Bioinformatics. 2011;27(12):1696–7.

62. Zhou Y, Zhou B, Pache L, Chang M, Khodabakhshi AH, Tanaseichuk O, et al. Metascape provides a biologist-oriented resource for the analysis of systems-level datasets. Nat Commun. 2019;10(1):1523.

