## Supplementary Figures for "Comprehensive epigenome characterization reveals diverse transcriptional regulation across human vascular endothelial cells"

**SUPPLEMENTAL FIGURES**

**Figure S1. Pairwise correlations of read distributions for all ChIP-seq datasets generated in this study.**

**Figure S2. Correlation heatmap of observed and expected gene expression values.**

**Figure S3. The distribution of active promoter and enhancer sites with respect to gene annotation information.**

**Figure S4. Read distribution at the RSPO3 locus.**

**Figure S5. Top four putative binding motifs identified from active promoters, distal enhancers and EC-specific enhancers.**

**Figure S6. Ratio of enhancer sites of each EC type that overlap with EC-specific enhancer sites.**

**Figure S7. GO terms (Biological Process) for active promoter sites, all enhancer sites and EC-specific enhancer sites.**

**Figure S8. PCA of gene expression data.**

**Figure S9. The relative log ratio of occurrence probability of DPs and DEs against DEGs.**

**Figure S10. k-Means clustering (k = 6) of the DP sites.**

**Figure S11. Hierarchical clustering heatmaps of log-scale gene expression data for claudin family members.**

**Figure S12. Boxplot of H3K27ac read density distribution among EC samples.**

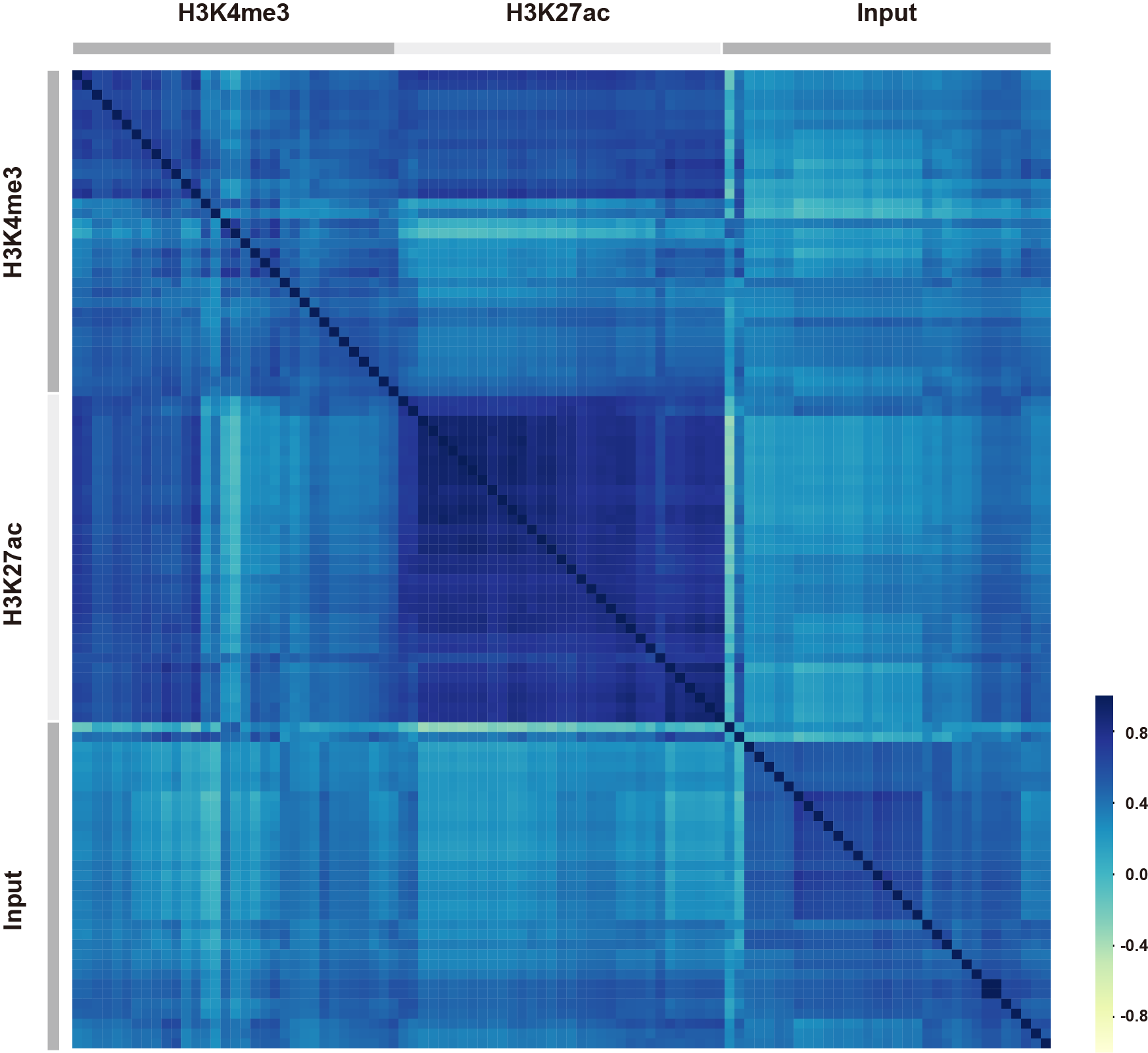

**Figure S1. Pairwise correlations of read distributions for all ChIP-seq datasets generated in this study.**

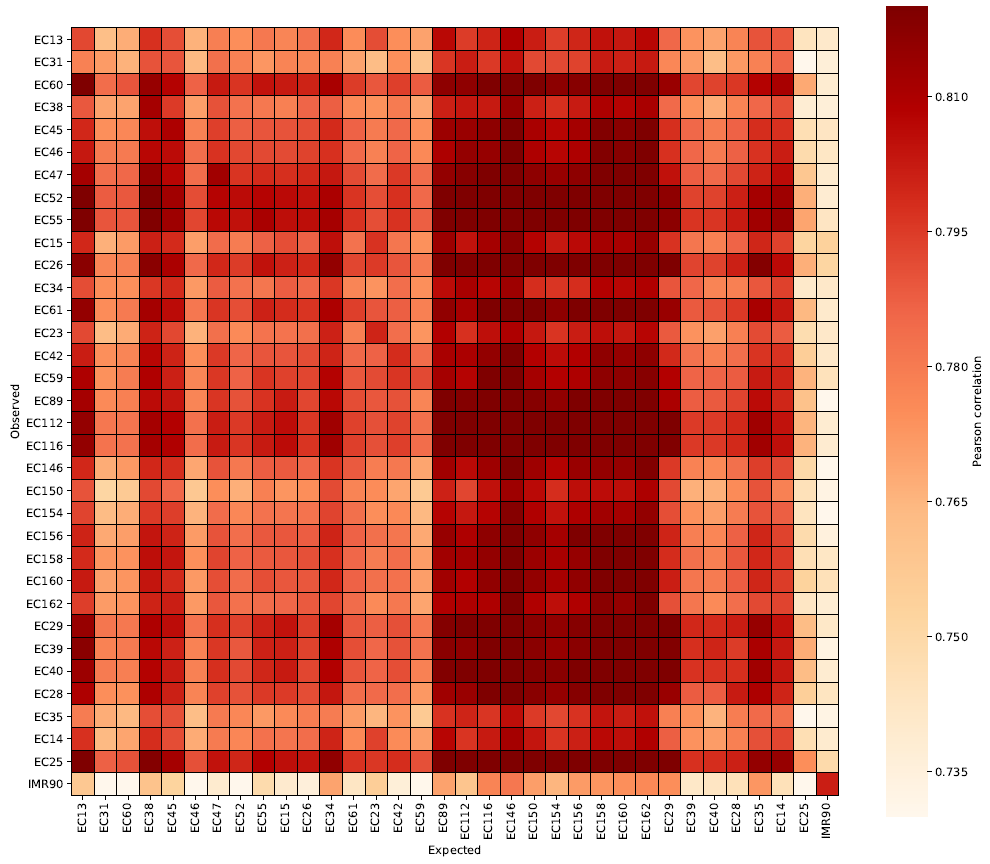

**Figure S2. Correlation heatmap of observed and expected gene expression values.**

(A) **Active promoter**

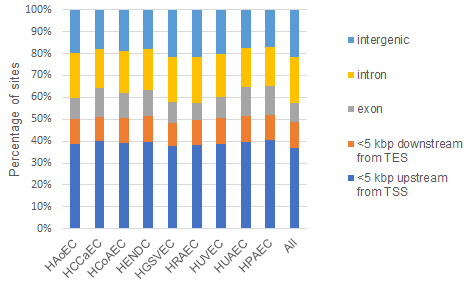

(B) **Enhancer**

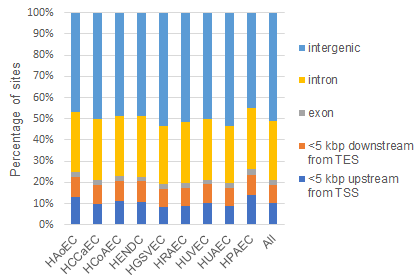

**Figure S3. The distribution of active promoter and enhancer sites with respect to gene annotation information.** (A, B) Data are shown for (A) active promoters and (B) enhancers. TES, Transcription End Site.

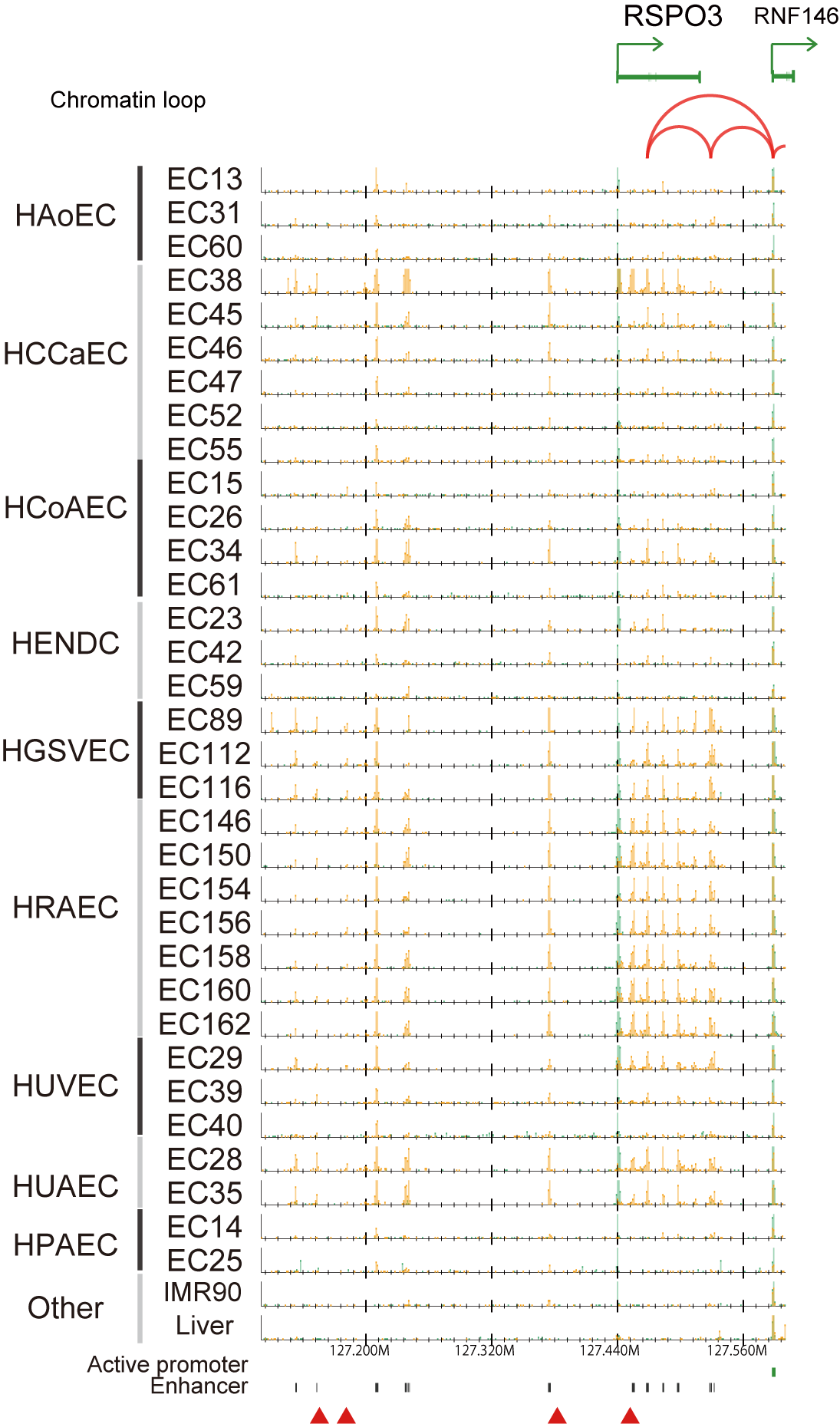

**Figure S4.** **Read distribution at the *RSPO3* locus.** Red triangles indicate GWAS SNPs obtained from GWAS Catalog.

Active promoter

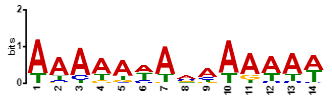

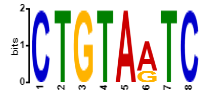

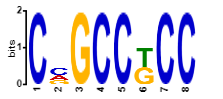

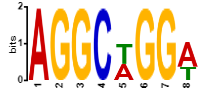

Distal enhancer

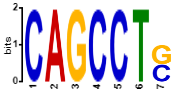

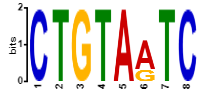

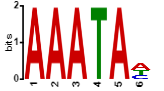

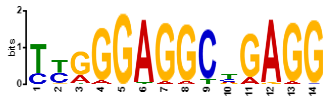

EC-specific enhancer

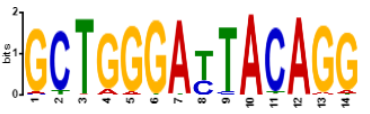

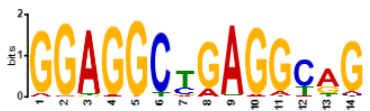

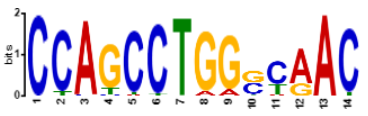

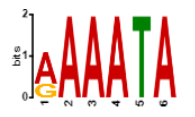

**Figure S5. Top four putative binding motifs identified from active promoters, distal enhancers and EC-specific enhancers.**

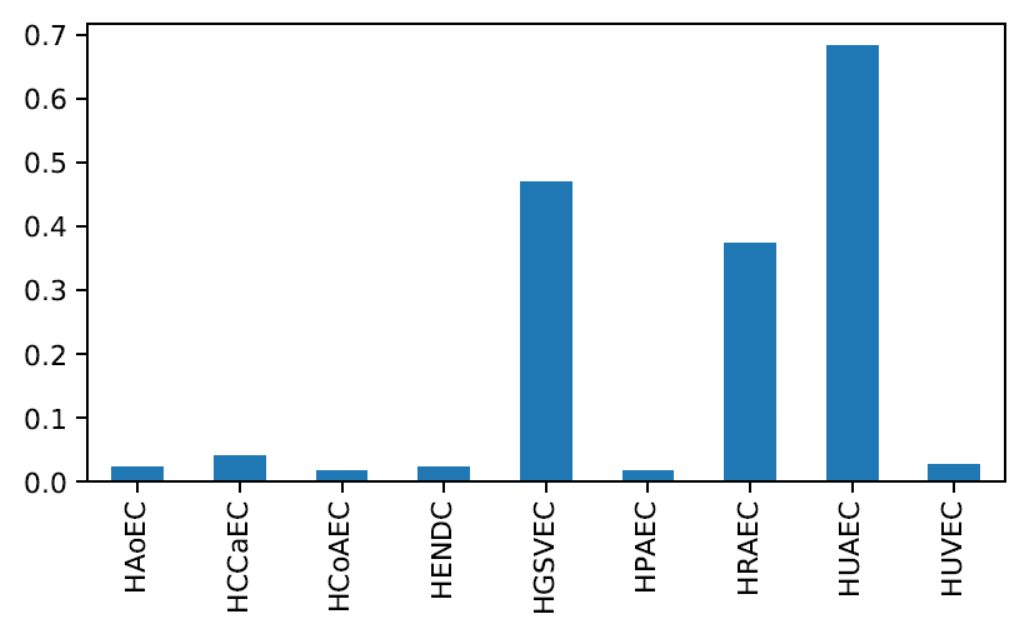

**Figure S6. Ratio of enhancer sites of each EC type that overlap with EC-specific enhancer sites.**

Active promoter

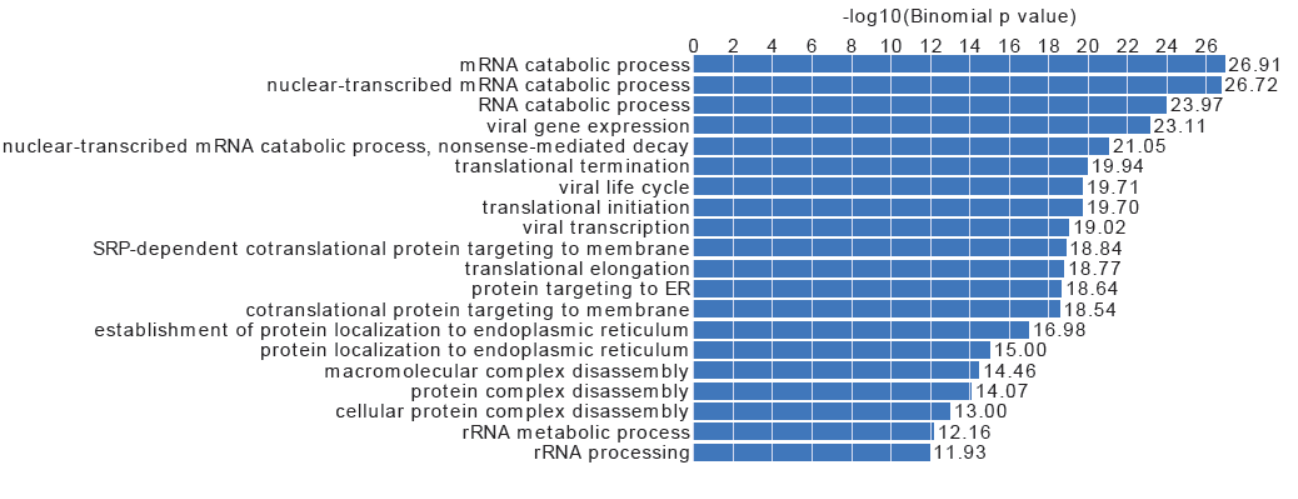

Enhancer

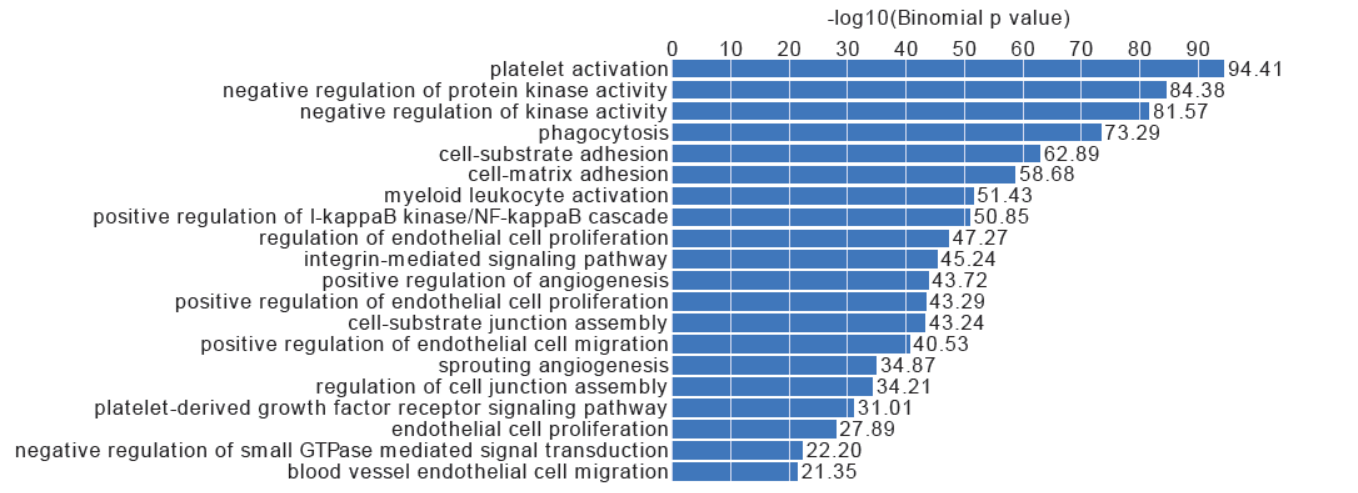

EC-specific enhancer

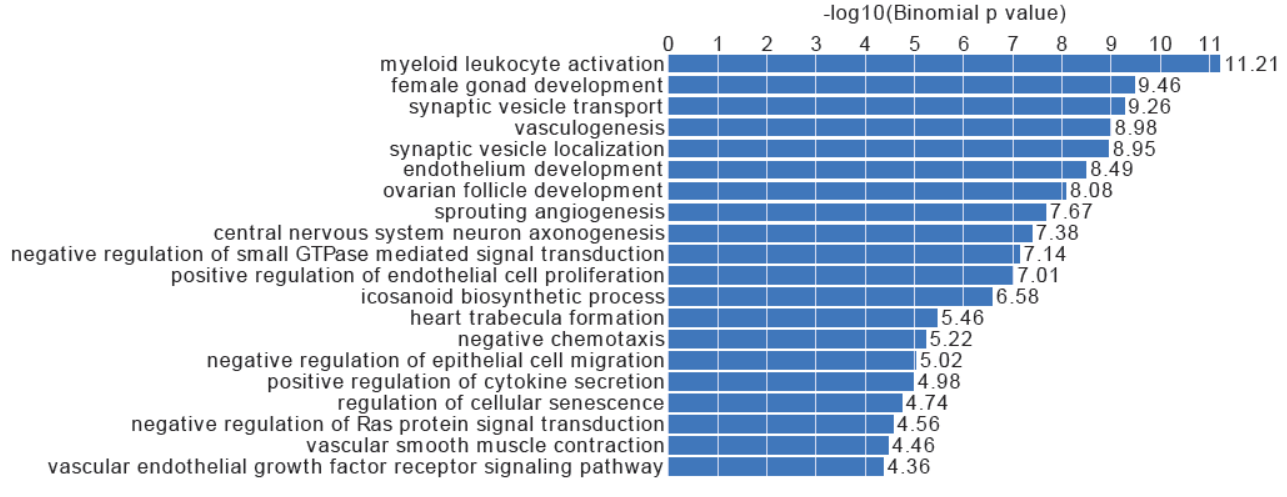

**Figure S7. GO terms (Biological Process) for active promoter sites, all enhancer sites and EC-specific enhancer sites.** These results were obtained by GREAT (<http://great.stanford.edu/public/html/>) with the default setting.

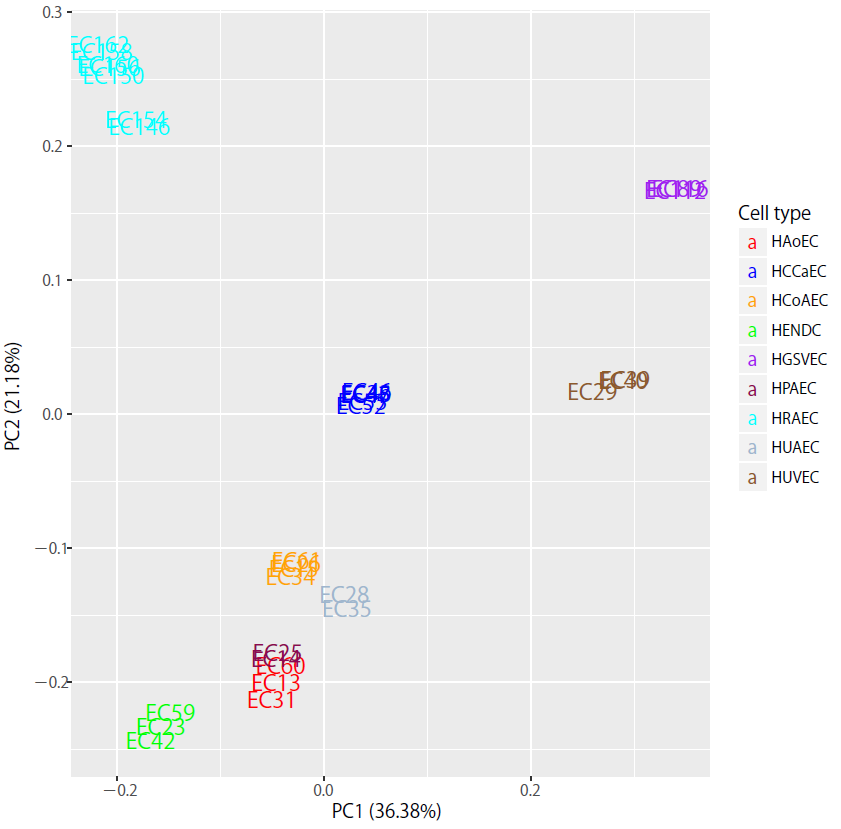

**Figure S8. PCA of gene expression data.** This figure is analogous to data shown in Fig. 4A for H3K27ac. The color of samples indicates EC types.

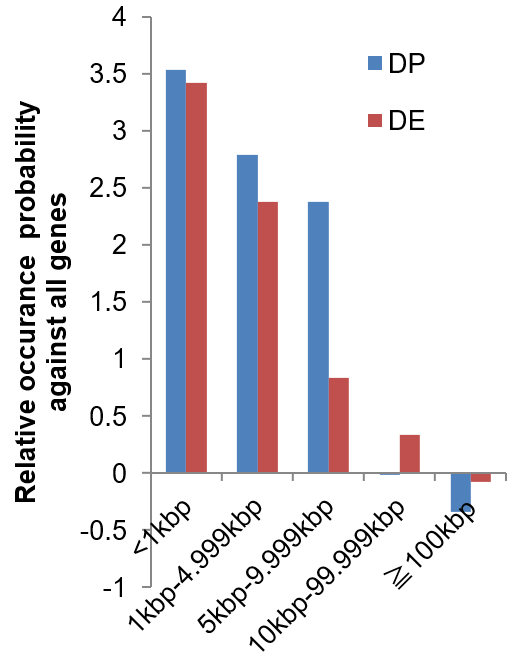

**Figure S9. The relative log ratio of occurrence probability of DPs and DEs against DEGs.** The *y* axis indicates the log ratio of the probability of occurrence of differential promoter sites (DPs, blue) and enhancer sites (DEs, red) around DEGs as compared with all genes. Positive values indicate that sites are more located within a distance from the TSS of DEGs compared with all TSSs as shown along the *x* axis.

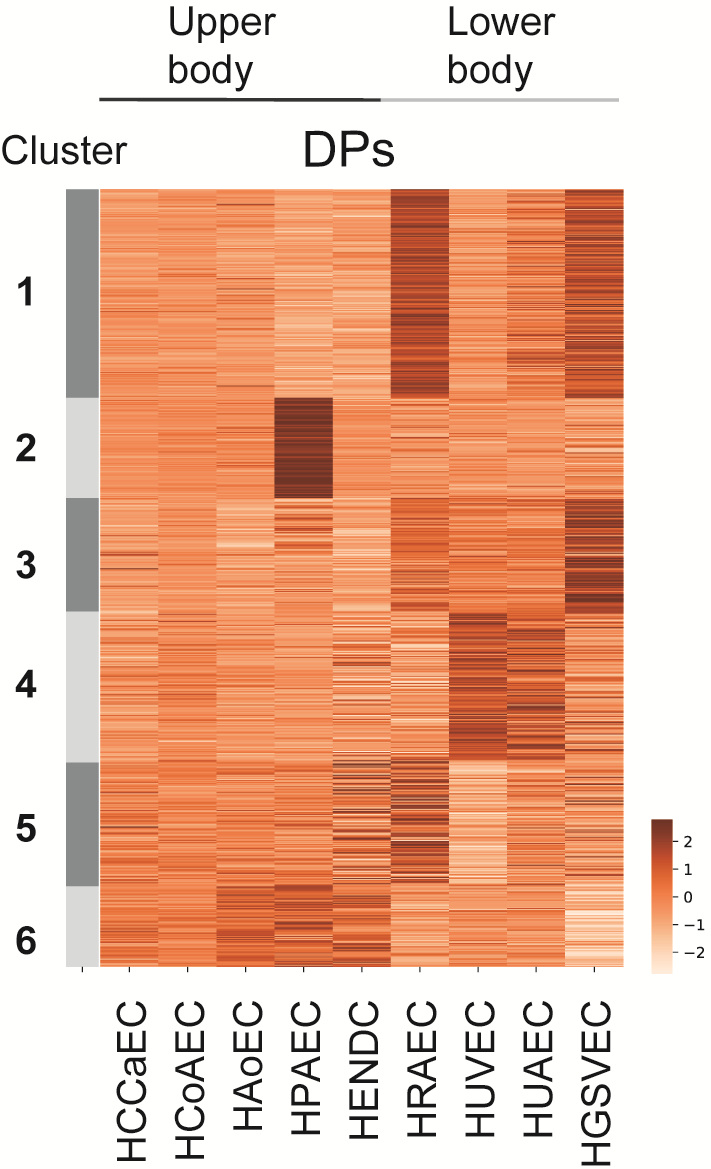

**Figure S10. k-Means clustering (k = 6) of the DP sites.** The color indicates the Z-scores across EC types (a representative for each type) for each site.

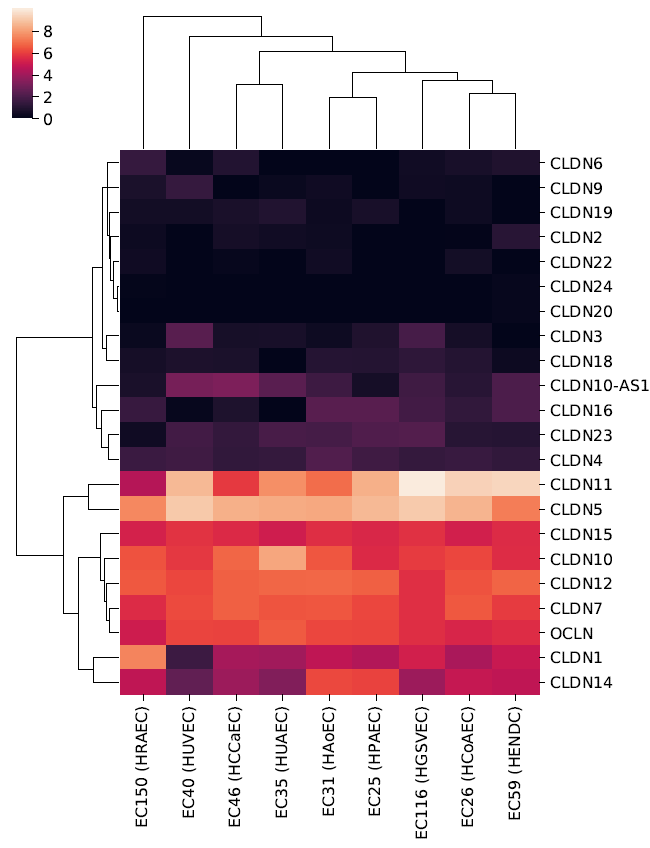

**Figure S11. Hierarchical clustering heatmaps of log-scale gene expression data for claudin family members.**

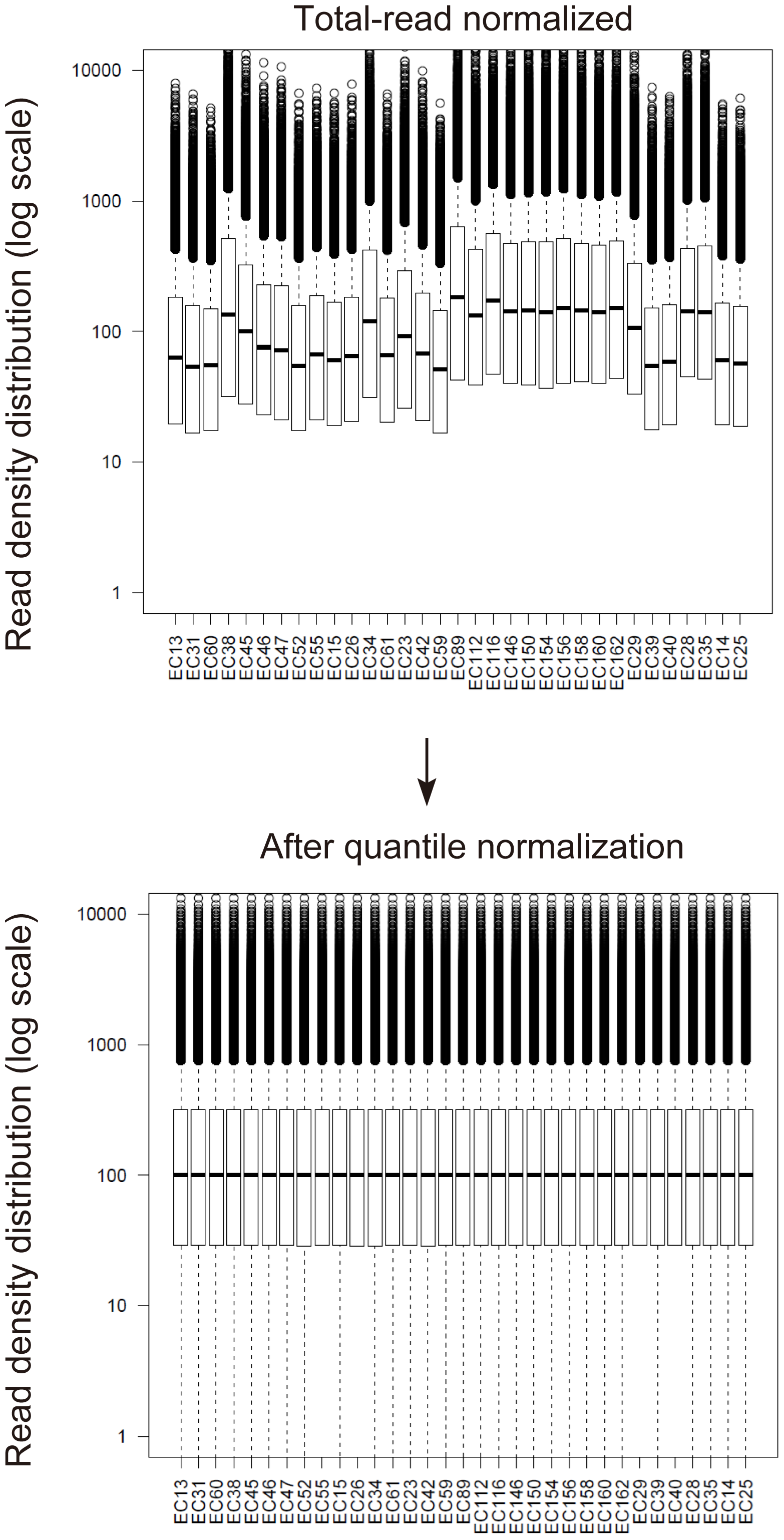

**Figure S12. Boxplot of H3K27ac read density distribution among EC samples.** The white box indicates first and third quartiles, and the black horizontal line indicates the median value. The upper graph shows the data normalized for total mapped reads. The lower graph shows data after quantile normalization.
